# DEC1 Regulates Human β Cell Functional Maturation and Circadian Rhythm

**DOI:** 10.1101/2025.04.03.647023

**Authors:** Sam Preza, Bliss Zheng, Zihan Gao, Akshaya Biju, Mai Liu, Zhihui Cheng, Matthew Choi, Juan R. Alvarez-Dominguez

## Abstract

Stem cell-derived islet (SC-islet) organoids offer hope for cell replacement therapy in diabetes, but their immature function remains a challenge. Mature islet function requires the β-cell circadian clock, yet how the clock regulates maturation is unclear. Here, we show that a circadian transcription factor specific to maturing SC-β cells, DEC1, regulates insulin responsiveness to glucose. SC-islet organoids form normally from *DEC1*-ablated human pluripotent stem cells, but their insulin release capacity and glucose threshold fail to increase during *in vitro* culture and upon transplant. This deficit reflects downregulation of maturity-linked effectors of glucose utilization and insulin exocytosis, blunting glycolytic and oxidative metabolism, and is rescued by increasing metabolic flux. Moreover, DEC1 is needed to boost SC-islet maturity by synchronizing circadian glucose-responsive insulin secretion rhythms and clock machinery. Thus, DEC1 links circadian rhythms to human β-cell maturation, highlighting the essential role of circadian control in generating fully functional SC-islet organoids.

## INTRODUCTION

The promise of broad cell therapy for diabetes has grown closer to reality with the advent of islet organoids grown from human pluripotent stem cells (SC-islets).^1–3^ Islet transplants can cure insulin-dependent diabetics^4^, but are limited by a scarcity of acceptable islets. SC-islets offer limitless cells for transplantation therapy, which can render human recipients free of daily insulin injections.^5,6^ Yet, *in vitro* SC-islets are molecularly and functionally less mature than transplanted ones^7–9^, and both lack the kinetics, precision, and magnitude of glucose-stimulated insulin secretion (GSIS) of adult islets.^10,11^ This delays therapeutic benefit from SC-islet transplants, posing a key challenge to their broad application.

Mature islet function develops after birth, as hormone secretion capacity and the glucose threshold for secretion increase.^12–14^ These changes are intertwined with the postnatal onset of islet circadian clocks.^15^ The clock in islet β cells aligns GSIS responsiveness with the active phase of the day^16^, likely to adapt to postnatal feeding-fasting rhythms.^8^ Inactivating CLOCK or BMAL1—the master clock activators—renders mouse islets unable to mount mature GSIS responses, causing diabetes.^17–20^ Whereas we find that inducing clock genes, via daily metabolic stimulation and recovery cycles, fosters mature function in human cadaveric/SC-islets by prompting circadian GSIS responses with a raised glucose threshold.^9^ How the clock coordinates metabolic maturation of human β cells, however, remains poorly understood.^10,21^

To elucidate the regulation of human SC-β cell maturation by the circadian network, we focused on the transcription factor DEC1 (also called BHLHE40, SHARP2). DEC1 entrains circadian rhythms to environmental cues, including light and feeding^22,23^, by competing with CLOCK:BMAL1 for DNA E-box binding.^24^ Recently, we found that mouse DEC1 coordinates islet GSIS by synchronizing energy metabolism and exocytic gene rhythms, and that *Dec1* knockout renders mouse islets immature, causing lifelong glucose intolerance due to insufficient insulin responses.^25^ We previously predicted that DEC1 partakes in the core regulatory circuit that defines human β-cell identity.^9^ However, *DEC1*’s role in human β cell development, functional maturation, and metabolism remains unexplored.

Here, we report that DEC1 is essential for SC-β cell maturation *in vitro* and *in vivo*. SC-islet organoids differentiate normally from *DEC1*-ablated human pluripotent stem cells (hPSCs) in scalable suspension culture^26^, but exhibit impaired GSIS in both static and dynamic assays. RNA-seq of purified SC-β cells links this defect to downregulation of maturity-associated genes for glycolysis, mitochondrial respiration, and insulin exocytosis. Accordingly, *DEC1*^-/-^ SC-islet organoids exhibit an energy deficit, prompted by lower glucose uptake, that hinders glycolysis and oxidative respiration and incites mitophagy. This renders *DEC1*^-/-^ organoids unable to mature even after transplantation into the kidney capsule of immunodeficient mice. We further find that DEC1 is required for enhancing SC-islet maturity through synchronization of circadian clock machinery and GSIS rhythms. Importantly, the GSIS deficit of *DEC1*^-/-^ organoids can be rescued pharmacologically, by insulinotropic agents that bypass metabolism or increase metabolic flux. These findings establish DEC1 as a key link between the circadian clock and human β-cell metabolic maturation, highlighting the potential of leveraging circadian control to generate SC-islets with adult-like GSIS for improved transplantation therapy.

## RESULTS

### DEC1 is required for mature insulin responses to glucose in human SC-islets

To evaluate DEC1’s role in human islet development, we tracked its expression at each stage of our scalable protocol for differentiating 3D SC-islet organoids in suspension bioreactors (Figure 1A).^26^ *DEC1* is minimally detectable in differentiating progenitors but enriched in SC-islet β cells, present in ∼97% of β cells and induced 12-fold (Figures 1B and S1F). We thus disrupted *DEC1* in independent hPSC lines using an inducible CRISPR-Cas9 system (Figure S1A).^27^ *DEC1^-/-^* lines differentiated normally, forming 3D SC-islet organoids with intact morphology, distribution of α (glucagon+/C-peptide+), β (C-peptide+/glucagon-≡ C-peptide+/NKX6.1+), and enterochromaffin-like (SLC18A1+/C-peptide-) cells, and insulin production (Figures S1C-S1G). We verified *DEC1* mRNA and protein loss throughout differentiation and *in vitro* maturation by extended culture (28 days) in serum-free media without added factors (Figures 1C, S1B, and S1F)^28,29^. DEC1 absence impaired induction of IAPP, a marker of SC-β cell functional maturation^30^ (Figures 1D and S1H). Strikingly, *DEC1^-/-^* SC-islets show markedly diminished GSIS responses in both serial static and dynamic assays (Figures 1E-G). Insulin secretion was significantly higher under static non-stimulatory (2.8mM) glucose incubations, while subsequent responses to saturating (20mM) glucose were impaired, despite normal insulin release following chemical depolarization with KCl (Figure 1E). Thus, *DEC1^-/-^* SC-islet organoids show elevated insulin secretion under non-stimulatory glucose and lower GSIS capacity—hallmarks of immature β-cell function.^12–14^

**Figure 1.**
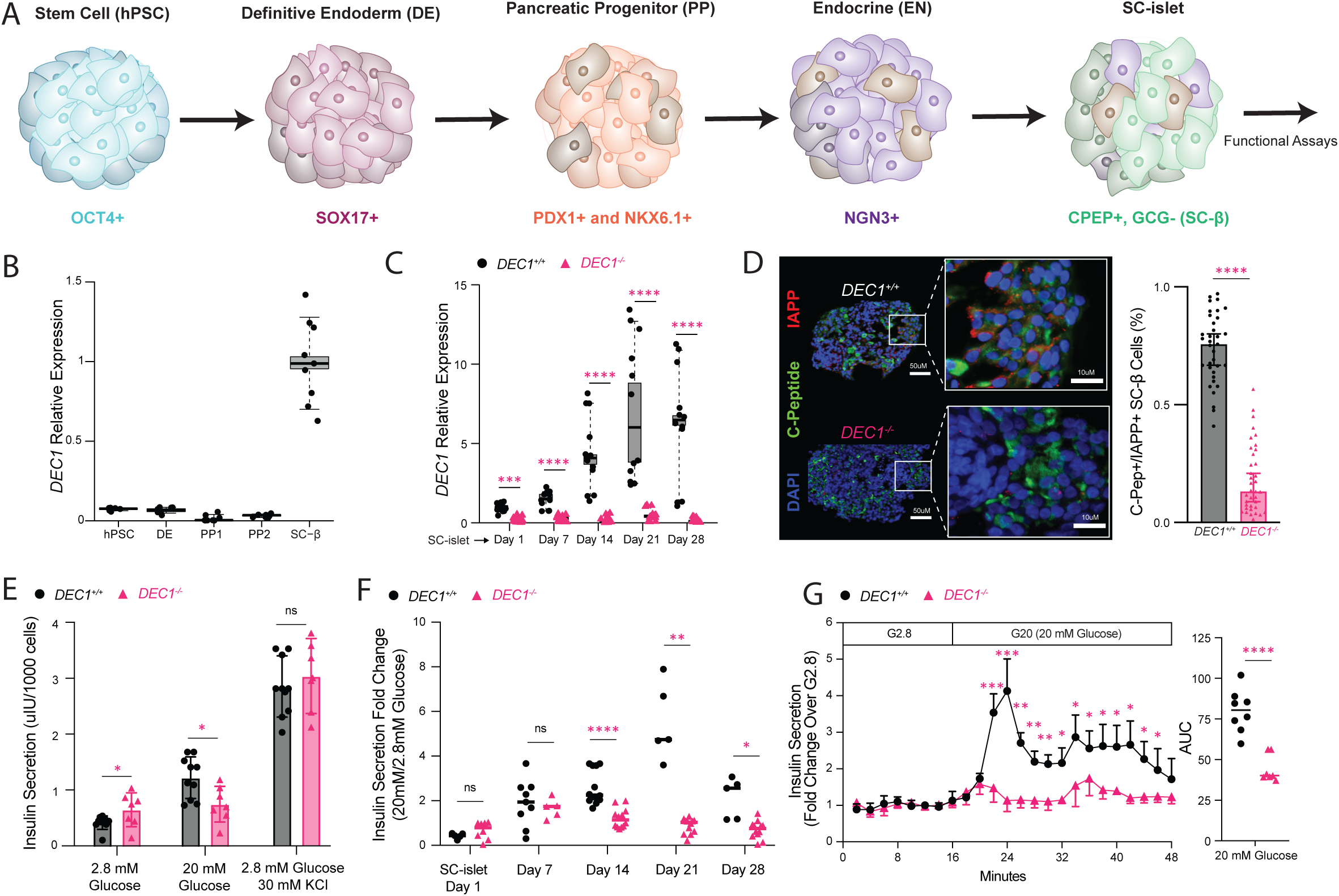
DEC1 is required for mature insulin responses to glucose in human SC-islets. (A) Stepwise directed differentiation of SC-islets from human pluripotent stem cells (hPSC). (B) *DEC1* expression is low through differentiation stages and selectively induced in SC-β cells. Data from N = 4 differentiations with n = 2 technical replicate measurements each, normalized to day 21 purified SC-β cells. (C) *DEC1* expression is abrogated in *DEC1^-/-^* SC-islets from week 1 through week 4 of extended *in vitro* culture. Data from N = 4 differentiations with n = 3 replicate technical measurements each, normalized to day 1 SC-islets. (D) *DEC1* loss impairs IAPP induction in SC-β cells. Images are C-peptide and IAPP immunostaining of day 14 *DEC1^+/+^* and *DEC1^-/-^* SC-islets (left). Percent of C-peptide positive cells that are IAPP positive across N=5 *DEC1^+/+^* and 5 *DEC1^-/-^* differentiations, quantifying n = 4-6 SC-islet organoids from each (right). Scale bar is 10 or 50um. (E) *DEC1*^-/-^ SC-islets show immature glucose stimulated insulin secretion (GSIS) responses under the indicated sequential static incubations, marked by elevated insulin secretion in non-stimulatory (2.8 mM) glucose, and decreased insulin secretion in stimulatory (20 mM) glucose. Data from day 14 *DEC1*^+/+^ SC-islets (N=5 differentiations, n = 50-100 SC-islets assayed from each) and day 14 *DEC1*^-/-^ SC-islets (N=4 differentiations, n = 50-100 SC-islets assayed from each), each assayed at least in duplicate. (F) *DEC1*^-/-^ SC-islets fail to expand GSIS responsiveness during extended *in vitro* culture. Data are insulin secretion fold-change between sequential static 2.8mM and 20mM glucose incubations of *DEC1*^+/+^ SC-islets (N=5 differentiations, n = 50-100 SC-islets each) and *DEC1*^-/-^ SC-islets (N=4 differentiations, n = 50-100 SC-islets each) at day 1, 7, 14, 21, and 28, each assayed at least in duplicate. (G) GSIS dynamics reveal *DEC1*^-/-^ SC-islets show low peak (first phase) insulin secretion under stimulatory (20 mM) glucose and dampened second phase secretion, consistent with an immature phenotype. Data from day 14 *DEC1*^+/+^ (N=3 differentiation, n = 50-100 SC-islets, each assayed at least in duplicate) and *DEC1*^-/-^ SC-islets (N=3 differentiations, n = 50-100 SC-islets, each assayed at least in duplicate) perifused with the indicated substrates, normalized to the mean of the first incubation. Area under the curve (AUC) summarized to the right. Data are mean ± SEM. *p <0.05, **p <1e-2, ***p <1e-3, ****p <1e-4, unpaired Welch’s t test [(E), (F), and (G)] and Wilcoxon test [(C) and (D)].

Longitudinal assays over four weeks of extended *in vitro* culture further revealed that *DEC1^-/-^* SC-islets fail to expand their GSIS capacity, as evidenced by consistently lower insulin secretion fold-change from 2.8mM to 20mM glucose (Figure 1F). In dynamic assays, although *DEC1^-/-^* SC-islets retain a biphasic pattern of insulin secretion, they show significantly reduced peak stimulation (4.14 ± 1.4-fold decrease) under saturating glucose, leading to diminished cumulative secretion (Figure 1G).

DEC1 both regulates and is regulated by the circadian clock.^22,31^ Given DEC1’s requirement for mature GSIS responses in SC-islets, we asked if the clock contributes to SC-islet maturity. To test this, we used siRNAs to deplete BMAL1 and PER2 in day 14 SC-islets, core transcriptional regulators of the clock’s activating and repressing arms, respectively (Figure S1I). *PER1* and *BMAL1* knockdown SC-islets showed diminished GSIS responses in serial static and dynamic assays, reminiscent of neonatal/functionally immature islets.

Collectively, our findings show that DEC1 is critical for refining GSIS responses during SC-islet maturation *in vitro*, including suppressing basal insulin secretion and expanding GSIS capacity.

### DEC1 regulates β-cell maturity-linked effectors of glucose metabolism and insulin secretion

To study DEC1’s molecular role in SC-islet maturation, we performed longitudinal RNA sequencing of *DEC1^+/+^* and *DEC1^-/-^*organoids at *in vitro* maturation days 7, 14, 21 and 28 (Figures S2A and S2B). 2,556 genes were differentially expressed, including suppression of *IAPP* and effectors of GSIS and metabolic signaling (*PCSK1*, *GLP1R*, and *SYT13*^32^) and induction of lipid metabolism regulators (*ACSL1*, *ELOVL2*). To investigate DEC1’s role specifically in SC-β cells, we performed RNA sequencing in CD49a-purified cells from both day 1 and day 14 SC-islets (Figures 2A, 2B, and S2C).^29^ We identified a total of 2,622 genes significantly affected by *DEC1* loss (Figures 2C-F and S2D-E). These comprise downregulated genes impacting β-cell function through roles as regulators or effectors of glucose import (*POU4F2*)^33^, sensing (*HK2*)^34^, glycolysis (*LDHB*)^35^, mitochondrial oxidative phosphorylation (*NDUFA4*)^36^, redox homeostasis (*TXNRD2*)^37,38^, and insulin secretion (*ADCY5*)^39^. Genes upregulated in *DEC1^-/-^* SC-β cells, by contrast, include liver genes (*ALB, AFP*); disallowed glycolytic enzymes (*LDHA, PDK3*); mitochondrial depolarization (*TSPO*, *RACK1*)^40,41^ and autophagy (*PINK1*, *MMP9*, *BNIP3*)^42–45^ machinery; and fatty acid metabolism enzymes (*ACSL1*, *ACSL6*, *ELOVL2*)^46–48^. Interestingly, core circadian clock genes (*BMAL1, CLOCK*, *PER2*) lack differential expression (Figure 2B). Lastly, we confirmed RNA-seq results by qPCR for a subset of genes (*LDHA, LDHB, SYT13, GCK*) in enriched SC-β cells (Figure 2E). Our qPCR validation showed that in addition to *IAPP,* the key maturity markers *MAFA* and *SIX2* are also depleted in *DEC1^-/-^*SC-β cells (Figure 2E). To assess this comprehensively, we examined a set of 2,218 genes differentially expressed across human β-cell maturation pseudotime^49^, and found that they are globally depleted, with *IAPP* and *PAX4* among the leading-edge subset of genes driving the downregulated β-cell maturity signature (Figures 2C-F).

**Figure 2.**
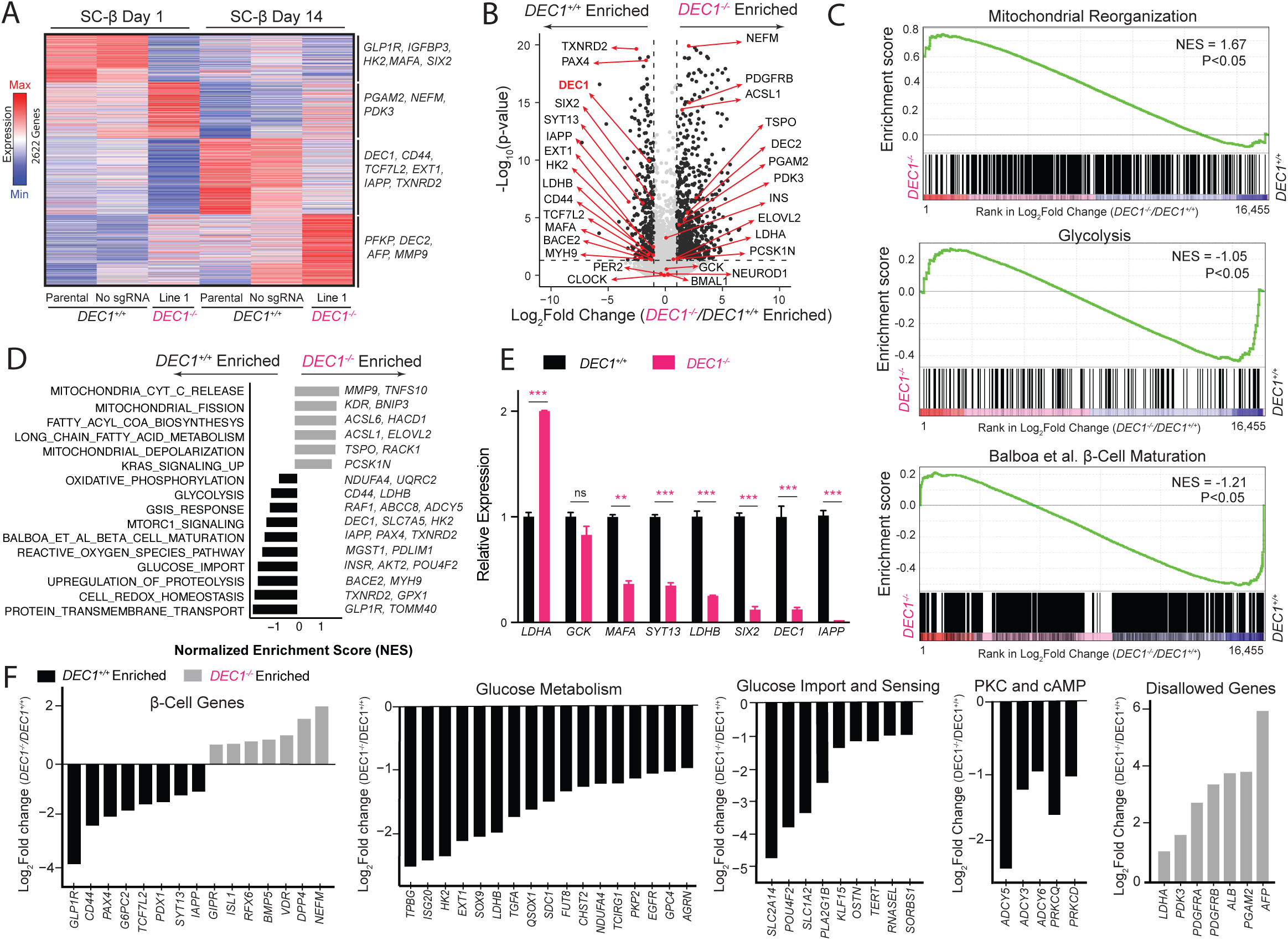
DEC1 regulates β-cell maturity-linked effectors of glucose metabolism and insulin secretion. (A) *DEC1* ablation in SC-β cells results in 2,622 differentially expressed genes. Heatmap shows row-normalized (z-scored) expression of differentially expressed genes (p < 0.05, DEseq test) in CD49a-sorted SC-β cells in Parental, No sgRNA, and *DEC1^-/-^* Line 1 at day 1 and day 14 of *in vitro* maturation. (B) SC-β maturity-linked genes are depleted in *DEC1^-/-^* SC-islets. Volcano plot shows average differential expression across SC-islet day 1, day 7 and day 14 for all differentially expressed genes at either day. Genes with ≥2-fold change and p-value < 0.05 are in black. Genes of interest are highlighted. (C) Mitochondrial reorganization genes are enriched in *DEC1^-/-^* SC-β, whereas glycolysis and β-cell maturity-linked genes are enriched in *DEC1^+/+^* SC-β cells. Gene set enrichment analysis shows genes ranked by fold change in *DEC1^-/-^*vs. *DEC1^+/+^* SC-β cells at day 14. NES, normalized enrichment score. (D) *DEC1* ablation disrupts induction of glucose stimulated insulin secretion (GSIS), glucose import, and redox homeostasis genes, and suppression of lipid metabolism genes, in SC-β cells. Gene set enrichment analysis shows pathways enriched in *DEC1^-/-^* vs. *DEC1^+/+^* SC-β cells at day 14. (E) Relative expression of selected genes from (B). Data are qPCR from day 21 SC-β cells. N = 3 differentiation, n = 2-3 technical replicates each. Data are mean ± SEM. *p <0.01, **p <1e-2, ***p <1e-3, Wilcoxon test. (F) *DEC1* ablation leads to upregulation of genes disallowed in mature β cells and downregulation of key glucose metabolism genes and insulin secretion genes. Shown are fold changes of genes in the indicated gene sets in day 14 SC-β cells.

Together, these findings implicate DEC1 in the control of GSIS maturation through the regulation of glucose import and metabolism, mitochondrial homeostasis, and insulin release.

### DEC1 controls circadian insulin secretion and clock gene rhythms in human SC-islets

The circadian clock attunes insulin secretion to daily feeding-fasting cycles^16^ and is needed for mature β-cell function.^16,50^ Previously, we reported that SC-islet organoids also display circadian GSIS responsiveness rhythms.^9^ Given DEC1’s ability to entrain circadian rhythms to environmental cues^22,23^, we investigated its requirement for GSIS rhythms. After synchronizing *DEC1^+/+^* and *DEC1^-/-^* SC-islets with 1-hour forskolin^20,51^ or dexamethasone^52^ pulses, we performed sequential static GSIS and collected RNA over two 24-hour cycles (Figure 3A). *DEC1^+/+^* SC-islets showed an autonomous 24-hour GSIS rhythm (p-value = 7.6E-07 from JTK and 9.70E-10 by RAIN analysis) first peaking at 36h post-synchronization (Figure 3B). At these peak times, GSIS capacity was on average 2-fold greater than in mock-treated SC-islets, which did not exhibit a GSIS rhythm (Figures 3B and 3C). This greater GSIS responsiveness was driven by lower insulin secretion under non-stimulatory (2.8mM) glucose (Figure S3A), as reported^9^. Strikingly, *DEC1^-/-^*SC-islets failed to show circadian GSIS responsiveness or suppressed basal insulin secretion upon synchronization, with no improvement in GSIS capacity (Figures 3B, 3C, and S3A). We verified circadian *DEC1* expression in synchronized but not mock-treated *DEC1^+/+^* SC-islets (Figure 3D). We also validated antiphasic expression of the core clock activator *BMAL1* and the *PER2* and *NR1D1* repressors in synchronized *DEC1^+/+^* SC-islets but not in mock-treated or *DEC1^-/-^* SC-islets (Figures 3D and S3B). Since *DEC1* regulates β-cell maturity and oxidative metabolism genes, we also examined expression patterns for *MAFA*, a key driver of mature β-cell^53,54^, and *DRP1,* which mediates mitochondrial^55^. Remarkably, DEC1 loss abrogates circadian *MAFA* and *DRP1* expression (Figure 3E).

**Figure 3.**
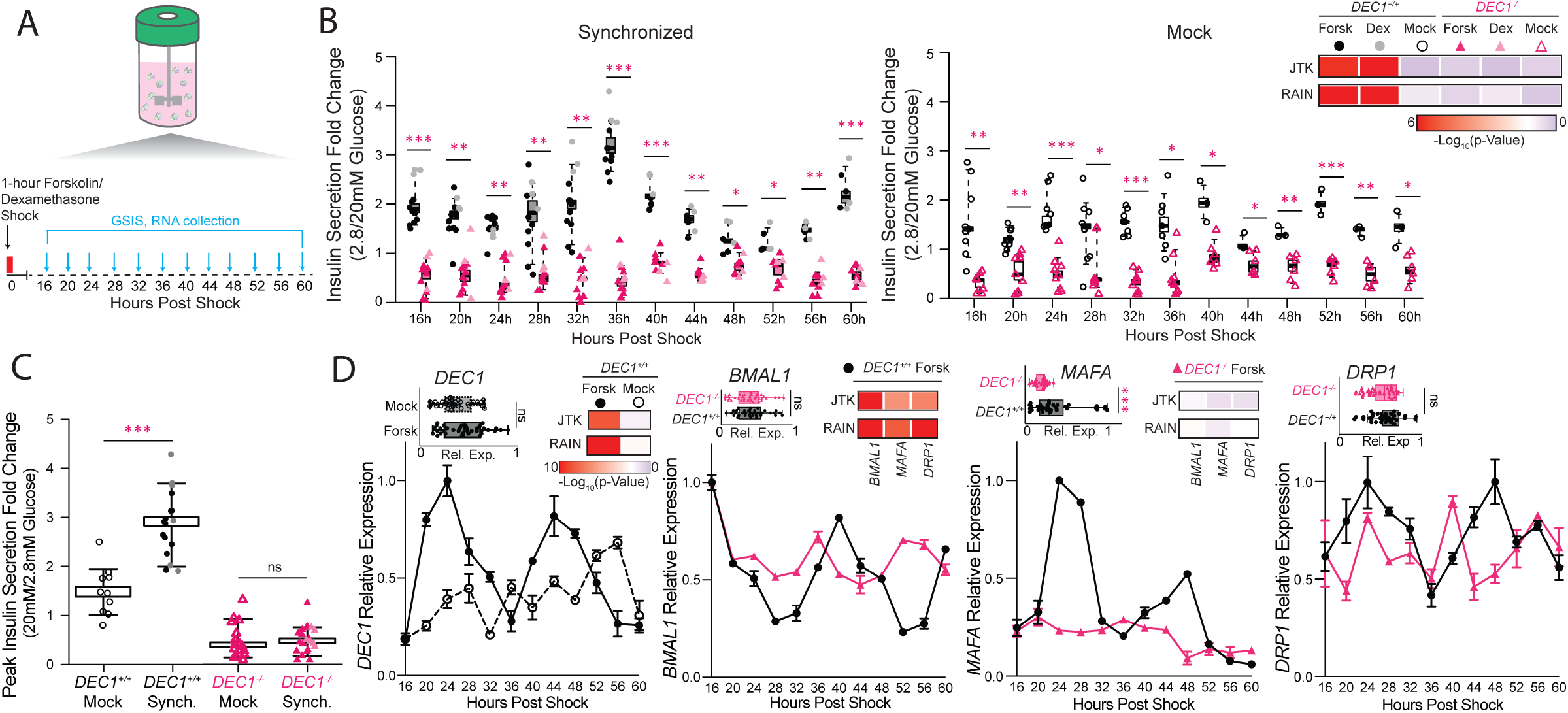
DEC1 controls circadian insulin secretion and gene expression rhythms in human SC-islets. (A) Experimental design for synchronization with 1-hour pulses of 10μM forskolin or 10μM dexamethasone. Glucose stimulated insulin secretion (GSIS) and RNA were assayed at 4 hour intervals 16 hours after synchronization. N=3 differentiations, with n=50-100 SC-islets assayed at least in duplicate. (B) *DEC1* is required for circadian SC-islet insulin secretion responses. Insulin secretion fold change follows a circadian rhythm (rhythmicity p-value < 0.05) in *DEC1^+/+^* vs. *DEC1^-/-^* day 14 SC-islets following forskolin or dexamethasone shock, with mock-treated controls shown to the right (N = 3 differentiation, n = 50-100 SC-islets). Shown are mean ± SEM insulin secretion fold changes at 16h post forskolin or dexamethasone shock and every 4h after. N=3 differentiations, with n=50-100 SC-islets assayed at least in duplicate. Rhythmicity p-values by JTK and RAIN analysis. (C) Insulin secretion fold-change is 2-fold greater in synchronized *DEC1^+/+^* SC-islets at the circadian peak time (36h or 60h post forskolin or dexamethasone shock) compared to mock controls. Data from (B). *p <0.05, unpaired Welch’s t-test. N=3 differentiations, with n=50-100 SC-islets assayed at least in duplicate. (D) Circadian and average *DEC1* expression following forskolin synchronization in *DEC1^+/+^* SC-islets (left). *DEC1* loss prevents circadian expression of *BMAL1, MAFA,* and *DRP1* following synchronization (rest). Data is expression relative to the maximum expression across the conditions shown in each panel (N = 3 differentiations with n = 2 replicate measurements each). Rhythmicity p-values by JTK and RAIN analysis.

Together, these data demonstrate that DEC1 is essential for proper circadian rhythms in GSIS and in the expression of core clock, mitochondrial dynamics, and β-cell maturity effectors.

### DEC1 sustains bioenergetics and mitochondrial integrity in maturing SC-β cells

Given DEC1’s role in regulating genes essential for respiration and redox homeostasis, we probed its influence on SC-β cell energetics and mitochondrial integrity. We find that DEC1^-/-^ SC-β cells show diminished mitochondrial membrane potential, along with elevated mitophagy (Figure 4A), consistent with mitochondrial dysfunction^45,56,57^. Analyzing oxygen consumption rates in day 14 *DEC1^-/-^* SC-islet organoids reveals severe reductions in basal, glucose-stimulated, ATP-linked, maximal, and non-mitochondrial respiration (Figures 4B, 4C, and S4A). Cellular reactive oxygen species levels are also significantly lower in *DEC1*^-/-^ SC-β cells (Figure 4A), in line with diminished respiration. Thus, DEC1 is vital to bolster mitochondrial oxidative activity driving glucose-coupled energetics.

**Figure 4.**
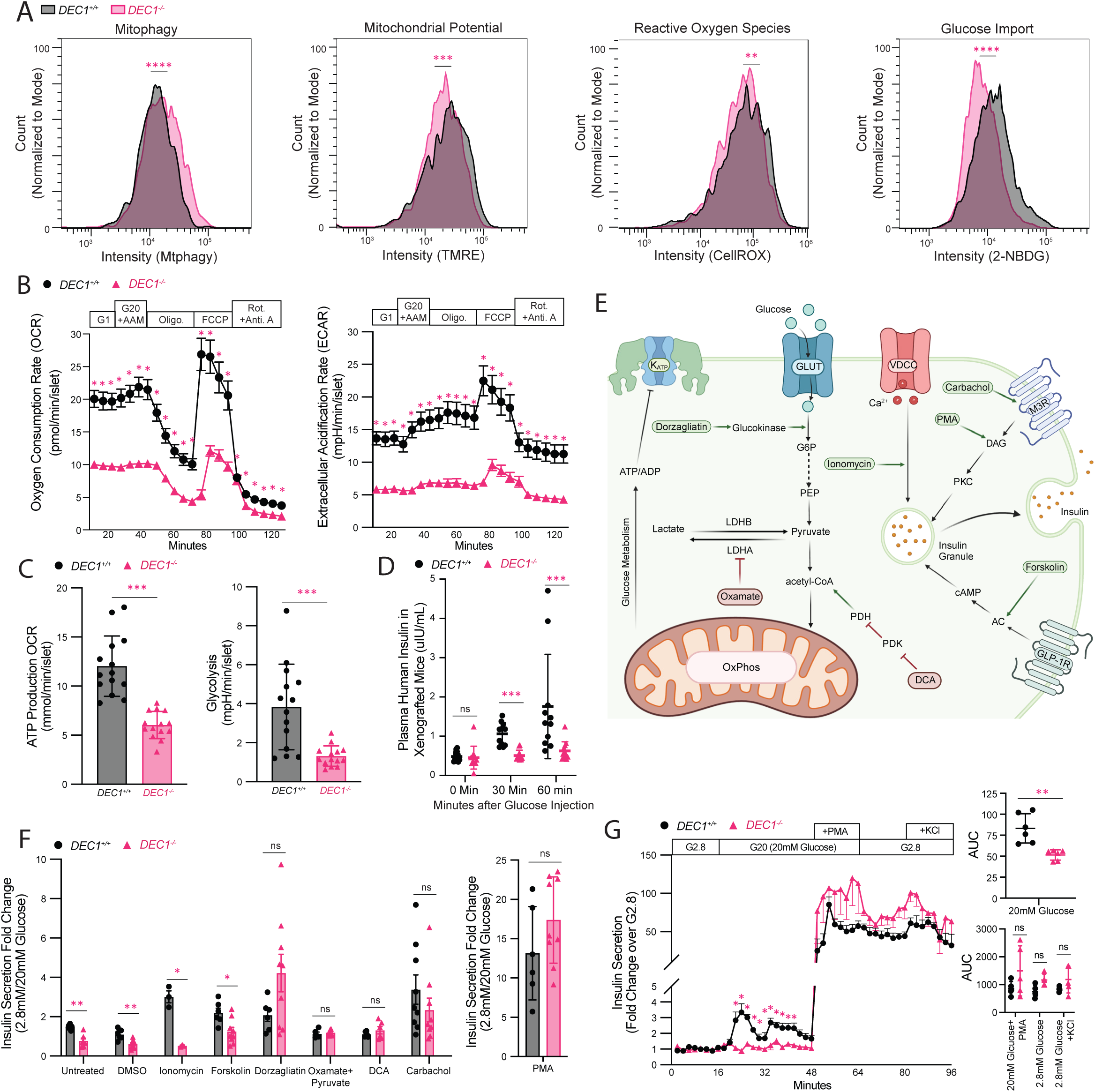
DEC1 sustains mitochondrial integrity and bioenergetics in maturing SC-β cells. **(A)** *DEC1*-ablated SC-β cells exhibit mitochondria with increased autophagy and reduced membrane potential, along with lower reactive oxygen species generation and glucose import. Data are distributions, normalized to mode, of flow cytometry for mitophagy (Mtphagy staining), mitochondrial membrane potential (TMRE staining), cellular reactive oxygen species (CellROX staining) and glucose uptake (2-NBDG staining) in CD49a-gated day 14 SC-β cells (N = 1 differentiation [2 more independent differentiations in Figure S4D], n ≥ 1000 cells). **(B)** *DEC1* loss stifles oxygen consumption rates (OCR, left) and extracellular acidification rates (ECAR, right). Data from day 14 *DEC1^+/+^* and *DEC1^-/-^* SC-islets in response to 20mM glucose and a physiological amino acid mixture (G20+AAM); oligomycin; FCCP; and rotenone with antimycin A. Data are mean ± SEM from N = 4 differentiations, each with n = 3-4 replicate measurements of 8 similar-sized day 14 SC-islets. **(C)** ATP production-linked respiration (left) and glycolysis (left) rates are impaired in *DEC1^-/-^* SC-islets. Data from (B). **(D)** Glucose stimulated insulin secretion defect in *DEC1*-knockout SC-islets is not rescued by *in vivo* kidney capsule transplant. Data are serum human insulin secreted into the blood of fasted immunodeficient mice transplanted with *DEC1^+/+^* or *DEC1^-/-^*SC-islets before, 30 minutes, and 60 minutes after intraperitoneal glucose injection 4 months post-transplant. Data are mean ± SEM from N = 4 differentiations with n = 11-12 mice each. **(E)** Pathway of glucose metabolism and downstream signaling targeted by insulinotropic agents screened for GSIS rescue experiments. **(F-G)** Glucose-independent PKC activator PMA, muscarinic receptor agonist carbachol, and metabolic flux enhancers (oxamate with methyl-pyruvate, DCA, and dorzagliatin) restore insulin secretion in day 14 *DEC1^-/-^* SC-islets under sequential static glucose incubations (F) (2.8 to 20mM glucose) or dynamic conditions for PMA (G) relative to *DEC1^+/+^* day 14 SC-islets. Data from N = 3 differentiations, each with n = 2-3 replicate measurements of 50-100 SC-islets. Data are mean ± SEM. *p <0.05, **p <1e-2, ***p <1e-3, ****p <1e-4, Wilcoxon test (A), and unpaired Welch’s t tests [(B), (C), (D), (F) and (G)].

Reduced mitochondrial activity upon DEC1 loss could reflect adaptations to reduced glucose utilization. Extracellular acidification rate analyses in day 14 SC-islet organoids reveal that glycolysis and glycolytic capacity are indeed hampered by *DEC1* loss (Figures 4B-C, S4A). Further, we trace such glucose utilization deficit to stifled glucose import, as shown by severely impaired uptake of a fluorescent glucose analogue in *DEC1^-/-^* SC-β cells (Figures 4A and S4D). Taken together with dulled glycolysis and mitochondrial oxidative activity^58^, these data argue that DEC1 promotes glucose import and utilization, stimulating the energy flux that underpins insulin secretion.

### A metabolic flux deficit underlies impaired insulin secretion upon DEC1 loss

To test the extent to which the maturation defect of *DEC1^-/-^*organoids can be rescued by the *in vivo* milieu, we transplanted day 21 SC-islets under the kidney capsule of immunocompromised mice. Strikingly, *DEC1^-/-^* SC-islets show blunted GSIS responses relative to *DEC1^+/+^* SC-islets even 19 weeks post-transplant (Figure 4D), as indicated by significantly reduced human insulin secretion following an intraperitoneal glucose injection. We thus conducted a targeted screen of insulinotropic agents to rescue the GSIS defect of *DEC1^-/-^* SC-islets *in vitro* (Figures 4E-G). Stimulation with forskolin, an adenylyl cyclase agonist, failed to restore GSIS, suggesting defects in steps upstream of cAMP-mediated secretory amplification. Ionomycin, a Ca^2+^ ionophore, also failed to rescue GSIS, consistent with normal Ca^2+^ influx dynamics in the absence of DEC1 (Figure S4B). Conversely, insulin secretion is restored to *DEC1^+/+^* levels by glucose-independent stimulation with either the diacylglycerol mimetic phorbol 12-myristate 13-acetate (PMA), or with carbachol, a muscarinic receptor agonist (Figures 4E-G). Thus, we sought to rescue the glucose coupling defect of insulin secretion in *DEC1^-/-^* organoids by increasing metabolic flux. Raising the pyruvate pool, via methyl-pyruvate supplementation combined with inhibition of lactate dehydrogenase A using oxamic acid, or increasing the acetyl-CoA pool, via inhibition of pyruvate dehydrogenase kinase with dichloroacetate (DCA), successfully restore GSIS to wild-type levels (Figure 4F). Similarly, enhancing glycolytic flux, via the glucokinase activator dorzagliatin, drastically improved GSIS to *DEC1^+/+^* levels (Figure 4F). Thus, enhancing metabolic flux rescues the GSIS defect of *DEC1^-/-^* SC-islets, demonstrating that DEC1 promotes glucose import and utilization to stimulate the coupling of insulin secretion to glucose metabolism in mature β cells.

## DISCUSSION

Transplantable SC-islet organoids are likely to become a therapeutic option for severe insulin-dependent diabetes. Yet, human SC-islets transplants today exhibit delayed therapeutic benefit, with 4 months needed for >85% of recipients to reach <7% HbA1c levels^6^, in contrast to 75 days or less for cadaveric islet transplants^59^. This delay likely reflects a need for SC-islets to complete functional maturation *in vivo*, in line with gradual molecular changes observed after transplant^9,60^. Common among these changes is the induction of circadian clock controllers^7,49^, and we previously showed that recreating circadian rhythms *in vitro* enhances SC-islet maturity^9^. How the circadian clock coordinates SC-islet maturation, however, has remained elusive.

Here, we show that a circadian transcriptional regulator, DEC1 (SHARP2, BHLHE40), is vital for metabolic maturation of SC-β cells *in vitro* and *in vivo*. We previously predicted DEC1 to be the most interconnected transcription factor in the core regulatory circuit defining mature human β-cell identity^9^. Consistently, DEC1 is specifically induced in maturing SC-β cells, and its absence thwarts induction of maturity-linked gene circuits. The disrupted circuits control glucose import and metabolism, mitochondrial dynamics, and insulin exocytosis. Accordingly, *DEC1*^-/-^ SC-islets show impaired glucose uptake, glycolysis, and oxidative metabolism. These changes stunt the ability of *DEC1*^-/-^ organoids to mount circadian GSIS responses, which fail to mature even after *in vivo* transplant, but are rescued by pharmacologically increasing metabolic flux. Thus, our findings demonstrate that DEC1 orchestrates the coupling of insulin secretion to glucose sensing, linking the clock to acquisition of the mature β cell phenotype.

Our report of a circadian islet maturation factor offers a new paradigm for how the clock programs metabolic specialization. Mature β-cell physiology is thought to be orchestrated by the CLOCK/BMAL1 complex, as inactivating *Clock* or *Bmal1* renders mouse islets immature^17–19,61^. However, roles for CLOCK/BMAL1 on human islet maturation are not well defined. Three lines of evidence argue that such regulation is indirect. First, CLOCK and BMAL1 are already present early in human β-cell development, before maturation^62^. Second, they are not part of the core regulatory circuit of mature human β cells^9^. Third, we find here that *DEC1* loss suffices to render SC-islets immature, despite intact *CLOCK* and *BMAL1*. Our data thus support DEC1 as the direct arm of the clock imparting circadian GSIS control to drive β-cell maturation.

Mechanistically, our results indicate that DEC1 mediates maturation in SC-β cells by coordinating glucose utilization downstream of core clock genes. DEC1-lacking β-cells exhibit reduced glucose influx and compensatory responses. These responses include induction of lactate dehydrogenase A, which diverts pyruvate away from mitochondrial metabolism toward lactate production, and of lipid utilization genes, likely signaling a shift to alternative energy substrates. These adaptations recall the metabolic flexibility of neonatal β-cells, which lack strict glucose-dependence for insulin release and show higher lipid utilization^10,14,63^. Notably, stimulating glucose utilization via glucokinase activation rescues GSIS to wild-type SC-β cell levels, ruling out glycolytic defects. Dorzagliatin stimulated insulin secretion more than inhibiting lactate dehydrogenase A or pyruvate dehydrogenase kinase—interventions expected to increase pyruvate and acetyl-CoA entry into the TCA cycle, respectively. These data provide further evidence for an intrinsic SC-islet bottleneck in glucose metabolism to pyruvate, occurring downstream of glucokinase but upstream of the TCA cycle, possibly through diversion into serine/glycine shunt.^49,64^

DEC1’s role in glucose metabolism is consistent with its ability to synchronize circadian rhythms with feeding, possibly through ChREBP-mediated nutrient sensing mechanisms, although these mechanisms are not well defined in islets^22,23^. Since islet circadian rhythms emerge postnatally^15^, along with the onset of daily feeding-fasting, they likely foster maturation, at least in part, through DEC1-mediated optimization of glucose metabolism to daily energetics. We further found, in separate mouse studies^25^, that DEC1 orchestrates mature GSIS responses by directly binding genes regulating integration of energy metabolism and insulin exocytosis to synchronize their expression, and thus energy and secretory rhythms. Taken together, our human and mouse studies reveal a new, evolutionarily conserved mechanism whereby DEC1 links circadian clockwork to β-cell energetics and metabolic specialization.

How can DEC1’s pathway be harnessed to improve SC-islet organoid maturity? Unlike global manipulation of metabolic cycles or the entire circadian network, precise tuning of DEC1 levels offers a new path to co-opt bioenergetic rhythms to control SC-β cell maturity. This may become possible by identifying DEC1’s nutrient sensing mechanisms that could be targeted for external modulation. The persistent functional defect of transplanted *DEC1*^-/-^ SC-islets suggests that such modulation would be relevant *in vivo*. Thus, further delineating how the circadian system underpins SC-β cell maturity heralds new avenues to attain fully functional SC-islets to improve islet replacement therapies.

### Limitations of Study

The experimental systems that were used in this study have limitations to consider. We used SC-islet organoids differentiated from the HUES 8 hPSC line, so genotype-specific effects were not evaluated. Our *in vitro* SC-islet differentiation protocol inherently produces some batch-to-batch differences in organoid functionality. Therefore, we assayed at least three independent SC-islet differentiations in each experiment. To assess *in vivo* function, we transplanted SC-islet organoids under the kidney capsule of immune deficient mice, so DEC1’s impact on SC-islet function under diabetic conditions was not investigated. Mechanistic links between DEC1 and target gene programs are inferred from perturbation studies; direct gene occupancy by DEC1 was not determined. Finally, our findings derive from an SC-islet organoid system that lacks the vascular, immune, and endocrine–neural inputs of native islets, so potential modulatory effects of blood perfusion, immune surveillance, and tissue crosstalk on DEC1-dependent maturation were not assessed.

## RESOURCE AVAILABILITY

### Lead Contact and Materials Availability

Further information and requests for resources and reagents should be directed to the Lead Contact, Juan R. Alvarez-Dominguez. All unique/stable reagents generated in this study are available from the Lead Contact with a completed Materials Transfer Agreement.

### Data and Code Availability

RNA-sequencing data generated in this study is publicly available at the Gene Expression Omnibus (GEO). The accession number for the raw and processed data reported is GSE302939.

## ACKNOWLEDGEMENTS

We thank Douglas A. Melton for support during the initial stage of this project; the University of Pennsylvania Diabetes Research Center Islet Cell Biology Core for technical support and critical discussions; and Israeli M. Galicia-Silva, Samuel D. Pollock, Niloo Rasouli, the University of Pennsylvania Stem Cell and Xenograft Core (SCXC – RRID:SCR_010035), the University of Pennsylvania Pathology and Imaging Core (RRID: SCR_022420), and members of the Alvarez-Dominguez laboratory for critical feedback on this manuscript. This work was supported by a grant to Sam Preza and J.R.A-D. from the Howard Hughes Medical Institute through the Gilliam Fellows Program; by NIH (K01DK129442), NIH/NIDDK DP1DK130673, NIH/NIGMS R35GM157320, the Human Islet Research Network (U24DK104162), start-up funding from the Perelman School of Medicine at the University of Pennsylvania, and a pilot award from the Diabetes Research Center at the University of Pennsylvania (P30DK19525) to J.R.A-D., who was also supported by a Howard Hughes Medical Institute Life Sciences Research Foundation Fellowship; and by grants to Douglas A. Melton from the JDRF (5-COE-2020-967-M-N), and the JPB Foundation (award no. 1094). We acknowledge using BioRender in the making of our graphical abstract (Created in BioRender. Preza, S. (2025) https://BioRender.com/864laoe).

## AUTHOR CONTRIBUTIONS

S.P., B.Z., Z.G., M.L., A.B., Z.C., M.C., and J.RA.-D. performed experiments. S.P., M.L., and A.B. conducted bioinformatics analyses. S.P., Z.G., and J.RA.-D. designed the research and interpreted results. S.P., B.Z., Z.G., A.B., and J.RA.-D. wrote the manuscript.

## STAR METHODS

### EXPERIMENTAL MODEL AND SUBJECT DETAILS

#### Human cell lines

The HUES8 (NIH hESC registry #09-0021; male) line was used for directed differentiation, and lines derived from HUES8 were used for CRISPR/Cas9 genome editing followed by directed differentiation. Undifferentiated cells were maintained as aggregates in supplemented mTeSR1 medium (StemCell Technologies) using spinner flasks (Corning) set at a 70rpm rotation rate in a 37°C 5%CO2 incubator.

### METHOD DETAILS

#### SC-islet differentiation in 3D suspension culture

Directed differentiation of human pluripotent stem cells into pancreatic islets was performed as previously described in 30-mL stirred suspension bioreactors (REPROCELL; ABBWVS03).^26^ All experiments were conducted using the HUES 8 (NIH registry #NIHhESC-09-0021) line maintained in mTeSR1 (StemCell Technologies, 85850) media on a rotator stir plate (Chemglass) at 70 RPM in a humidified 37°C, 5% CO₂ tissue culture incubator. Maintenance cultures were passaged every 3 days using a sequential filtration approach: 300 μm strainers were used to remove large aggregates, followed by 37 μm strainers (StemCell Technologies, 27250) to eliminate small debris. Cells were then incubated with Gentle Cell Dissociation Reagent (GCDR, StemCell Technologies, 100-0485) for 6 minutes, counted for viability, and reseeded at 0.6 × 10⁶ cells/mL in mTeSR1 supplemented with 10 μM ROCK inhibitor (Y27632, Abcam). Cells underwent at least three passages after thaw from cryopreservation before differentiation, with each passage achieving a minimum threefold expansion. A six-stage differentiation protocol was used to generate SC-islets. Cells were seeded at 0.6 × 10⁶ cells/mL in mTeSR1 with Y27632 on day 0, typically using 18M cells in 30 mL total volume. A half-media change was performed at 24 hours, replacing with fresh mTeSR1 (without Y27632), followed by a full media change on day 2. Differentiation was initiated on day 3 (Stage 1) and proceeded through Stage 6, with media changes performed on a 24-hour feeding cycle as described^26^.

#### Reverse transcription-quantitative PCR (RT-qPCR)

RNA extraction and purification was done using the DirectZol RNA miniPrep (Zymo Research, R2050), which included a DNase treatment to eliminate genomic DNA contamination. Complementary DNA (cDNA) synthesis was carried out using the High-Capacity cDNA reverse transcription kit (Invitrogen, 18080051), according to manufacturer instructions. qPCR reactions were performed using SYBR green master mix (Applied biosystems, A25742) on a StepOnePlus Real-Time PCR system (Applied biosystems). All reactions were done in triplicate, and melting curves were run to establish specificity of amplification. The primer sequences used for amplification are listed in the Resources Table. All qPCR CT values are normalized to 18S rRNA (ΔCT).

#### Static glucose-stimulated insulin secretion

Static glucose-stimulated insulin secretion (GSIS) assays were performed on Stage 6 clusters^26^. Clusters were first washed and equilibrated in KREBs buffer (128 mM NaCl, 5 mM KCl, 2.7 mM CaCl2, 1.2 mM MgSO4, 1 mM Na2HPO4, 1.2 mM KH2PO4, 5 mM NaHCO3, 10 mM HEPES, 0.1% BSA) containing 2.8 mM glucose for 1 hour. Then, we used 12 μm polycarbonate Millicell inserts (Sigma-Aldrich, PIXP01250) in 24-well plates to sequentially incubate clusters in low glucose (2.8 mM, 1 hour), high glucose (20 mM, 1 hour), and KCl (30 mM in 2.8 mM glucose, 1 hour). Media samples were collected after each incubation period. Following the assay, clusters were dispersed into single cells using TrypLE (Fisher Scientific, 50-591-419), which were counted to normalize secretion data. Insulin concentrations in the collected samples were measured using an ultrasensitive human insulin ELISA (ALPCO, 80-INSHUU-E01.1), with results expressed as μIU/mL/1000 cells. For insulin content measurements, cluster lysates were prepared using M-PER (Thermo Fisher, 78501) and analyzed using the same ELISA protocol. All assays were performed in triplicate.

#### Dynamic glucose-stimulated insulin secretion

Dynamic glucose-stimulated insulin secretion assays were conducted using an automated perifusion system (BioRep). Size-matched islets were carefully selected and placed in perifusion chambers. The chambers were first equilibrated with KREBs buffer containing basal glucose (2.8 mM) for 60 minutes at a constant flow rate of 100 μL/min. Following equilibration, islets were sequentially exposed to low glucose (2.8 mM) for 16 minutes, then high glucose (20 mM) for 32 minutes. For dynamic GSIS with insulinotropic agent, (10µM) PMA an additional 20 mM glucose step incorporating PMA was done for 16 minutes, followed by 16 minutes 2.8mM glucose, and 16 minutes of 2.8 mM glucose + 30 mM KCl. Insulin secretion data were normalized to the mean insulin release during the initial basal incubation.

#### Flow cytometry for differentiation cell markers

Suspension culture Samples were treated with TrypLE to dissociate clusters to single cells, which were then fixed in 4% Paraformaldehyde for 30 minutes at 4°C. The fixed cells were then blocked at 4°C for 30 minutes in a blocking buffer (5% Donkey serum [Jackson Immunoresearch], 0.1% Saponin [Sigma Aldrich] in PBS) to permeabilize cell membranes and minimize nonspecific binding. Primary antibody incubations were conducted overnight at 4°C, with antibodies selected based on the differentiation stage of the cells. The following day samples were incubated at room temperature for 1 hour with the appropriate fluorophore-conjugated secondary antibodies. The cells were resuspended in a washing buffer (5% donkey serum, PBS) and analyzed using an Accuri C6 Plus flow cytometer (BD Biosciences). Data acquisition and analysis was performed using FlowJo software. The antibodies used are listed in Resource Table and are as previously reported.^26^

#### Immunofluorescence Staining

Measurements were performed based on previously established protocols.^9,26^ SC-islet organoids were collected and fixed in 4% paraformaldehyde for 15 minutes at room temperature and washed with PBS. After fixation organoids were resuspended in embedding gel (2.5% agar, 2.5% gelatin). A droplet of the solution was transferred into an embedding cassette, stored in 70% ethanol, and then submitted to the University of Pennsylvania Pathology and Imaging Core (RRID: SCR_022420) for paraffin embedding. For deparaffinization the slides were treated with xylene, followed by serial ethanol and water washes. Antigen retrieval was done by incubating the slides in warm 1X Citrate buffer (Sigma Aldrich, C9999-1000ML) using a crockpot water bath at “high” temperature for 5 minutes and then “warm” for 30 minutes. The slides were subsequently blocked and permeabilized using a blocking buffer consisting of 5% donkey serum, 0.3% Triton and PBS solution for 1 hour to minimize nonspecific binding. Primary antibody staining was conducted overnight in 4°C for C-PEPTIDE, DEC1, Glucagon and SLC18A1 (Resource Table). After incubation with primary antibodies the slides were incubated with the appropriate secondary antibodies for an hour at room temperature and then counterstained with DAPI (Thermo Fisher, P36935). After staining, fluorescent imaging was performed using a fluorescent microscope.

#### SC-β cell enrichment with CD49a

Fluorescence-activated cell sorting (FACS) was performed as previously reported^29^ to isolate live, marker-defined β-cell populations from Stage 6 cultures. Cells were first dissociated into a single-cell suspension using TrypLE enzymatic digestion. Aggregates were dispersed mechanically through repeated pipetting with a P1000 micropipette, ensuring uniform suspension while minimizing cellular stress. The digestion reaction was quenched with S3 medium, followed by centrifugation at 300 × g for 5 minutes. The pellet was resuspended in sorting buffer (PBS + 1% BSA + 2 mM EDTA), filtered through a 35 µm strainer, and counted to determine cell concentration. For immunostaining, cells were aliquoted at 10 × 10⁶ cells/mL and incubated with anti-human CD49a-PE (1:11 dilution) (Miltenyi Biotec, 130-101-397) antibody for 20 minutes at room temperature, protected from light with gentle agitation every 3–4 minutes. After incubation, cells were washed twice by centrifugation (300 × g, 5 minutes) and resuspended in sorting buffer at a final concentration of 10 × 10⁶ cells/mL. FACS sorting was performed using a 100 µm nozzle at a rate of ∼10,000 events per second. CD49a⁺ cells were selectively sorted to enrich for insulin-producing β-cells within the Stage 6 population. Cells were collected into FACS tubes containing 1 mL sorting buffer supplemented with 1:1000 ROCK inhibitor (Y27632) to enhance post-sort survival. Sorted β-cells were kept on ice and subsequently transferred to pre-warmed media for further functional characterization or RNA extraction for downstream RNA-sequencing.

#### RNA-sequencing and differential gene expression

RNA-sequencing was performed as previously described.^9^ RNA from CD49a enriched SC-β cells (∼300k cells) or SC-islets (∼300k cells) was isolated for RNA-sequencing. Sequenced reads were STAR^65^ aligned using default parameters. Read counts per gene were quantified using HTSeq^66^ and normalized using DESeq.^67^ The expression levels were normalized to counts per million (CPM), with genes considered expressed if CPM was greater than 1.

#### Gene set enrichment analysis

Gene lists ranked by log_2_fold change (*DEC1^-/-^/DEC1^+/+^*) were analyzed for enrichment of genes using GSEA^68^ with default parameters and “-metric log2_Ratio_of_Classes.” Pathway annotations were derived from Hallmark and C5 (Gene Ontology) databases to investigate the overrepresentation of curated pathways and biological processes.

#### Forskolin and Dexamethasone synchronization experiments

Forskolin and Dexamethasone was used to synchronize circadian rhythms^20,51^ prior to downstream assays. Stage 6 SC-islet organoids were incubated in Stage 3 media supplemented with 10µM forskolin (Stemgent, 04-0025) or 10µM dexamethasone (Sigma, D4902-500mg) for 1 hour. Following the 1-hour forskolin or dexamethasone shock, SC-islets were returned to culture and allowed to recover in fresh Stage 3 media for an additional 15 hours. After the 15-hour post-shock incubation period, SC-islets were harvested for downstream analyses including static GSIS and RT-qPCR. Insulin secretion fold changes and relative expression of genes of interest were analyzed for circadian rhythmicity using both the JTK_CYCLE and RAIN algorithms implemented in Nitecap61.^69^ JTK_CYCLE employs nonparametric rank-based testing to detect periodic signals by comparing time-series data against idealized symmetric waveforms and assesses rhythmicity significance by fitting the data across multiple candidate periods. This process yields empirical p-values and estimates of the peak phase within a 24-hour cycle.

In parallel, RAIN (Rhythmicity Analysis Incorporating Nonparametric Methods) was applied as a complementary nonparametric approach. It tests rhythmicity by evaluating whether the data exhibit a significant rising trend followed by a falling trend within each cycle, allowing the peak to occur at any phase. The method uses exact null distributions derived from randomization principles to compute p-values, offering enhanced robustness to asymmetric waveforms and outliers. Both methods were used to test a 24-hour period.

The MetaCycle meta2d() function was used with default parameters to calculate rhythmicity metrics, including Benjamini-Hochberg corrected p-values and phase estimates. For visualization, time-series data was normalized to max expression.

#### Knockdown of *PER1* and *BMAL1* Using siRNA

SC-islets at day-14 were cultured in 6-well ultra-low attachment plates at a density of 1×10^6^ cells per well in complete medium. For each well, 9 µL of Lipofectamine® RNAiMAX was diluted in 150 µL of Opti-MEM® I Reduced-Serum Medium (Thermo Fisher Scientific). Separately, 3 µL of siRNA (20 µM stock; see resource table for product number) was diluted in 150 µL of Opti-MEM® Medium. The diluted siRNA was combined with the diluted Lipofectamine® RNAiMAX reagent at a 1:1 ratio (v/v), mixed gently, and incubated for 5 minutes at room temperature to allow complex formation. The resulting siRNA-lipid complexes (300 µL total volume) were added dropwise to each well.

Following transfection, plates were placed on an orbital shaker rotating at 300 rpm in incubator for 72 hours. The knockdown efficiency was validated via quantitative reverse transcription polymerase chain reaction (qRT-PCR). Functional assessment of insulin secretion was subsequently performed using static and dynamic GSIS assay.

#### Cellular and mitochondrial energetics analysis

Flow cytometry analyses of mitochondrial autophagy, mitochondrial membrane potential, cellular reactive oxygen species, and glucose analogue uptake were conducted in day 14 SC-islet organoids. Clusters were dissociated into single-cell suspensions using TrypLE and resuspended in Stage 3 media. Single-cell suspensions were incubated with a freshly prepared staining solution containing 50nM TMRE (Abcam; ab113852), 5µM CellROX Deep Red (Invitrogen, C10422), 20µM 2-NBDG (Thermo Fisher, N13195), or 100nM Mtphagy dye (Dojindo, Mt048) in Stage 3 medium. CD49a antibody (1:50) was included during incubation to gate-select for SC-β cells. Cells were incubated at 37°C for 30 minutes in a humidified 5% CO₂ incubator. After staining, cells were washed twice with 1% BSA in PBS and filtered through a 35 µm nylon mesh into flow cytometry tubes (STELLAR SCIENTIFIC, FSC-9005). Flow cytometry data was acquired using a BD Accuri C6 Plus flow cytometer (BD Biosciences) and analyzed using FlowJo v10.8.1 (BD Life Sciences). Fluorescence intensities were normalized to unstained controls and data were gated based on forward and side cell scatter parameters to exclude debris and doublets.

#### Metabolic flux analysis

Oxygen consumption rate (OCR) measurements were performed using the Seahorse XFe96 Analyzer and Spheroid FluxPak (Agilent, 102905-100) to assess mitochondrial function in SC-islets at day 14 of *in vitro* maturation. One day prior to the assay, sensor cartridge wells were hydrated by incubating with 200 µL of Seahorse calibrant solution (Agilent) at 37°C overnight in a non-CO₂ incubator. Spheroid plate wells were coated with 30 µL of 100 ng/mL poly-D-lysine (Sigma, P7280). Injection basal media (Agilent; 103335-100), wash media (basal media with 0.5% FBS), and assay media (basal media with 0.5% FBS and 1mM glucose) were freshly prepared and stored at 4°C protected from light. On the day of the experiment, all media were adjusted to pH 7.4 and warmed and kept at 37°C. SC-islets were hand-selected and washed twice in wash for 15 minutes at room temperature. After washing, 8-10 size-matched SC-islets were transferred per well in a total volume of 30 µL using an islet-loading mold and gently overlaid with 175 µL of pre-warmed assay media. The Spheroid plate was incubated for 1 hour at 37°C in a CO₂-free incubator to allow for metabolic equilibration. Prior to the assay, the sensor cartridge was calibrated on the Seahorse XFe96 Analyzer for 15 minutes. Mitochondrial stress testing was then performed by sequential injection of 20 µL the following compounds, prepared in injection basal media and loaded into the cartridge injection ports: 20 mM glucose with 0.5mM glutamine and 3.5mM amino acid mix, 5 µM oligomycin (Sigma, O4876), 2 µM FCCP (Sigma, C2920), and a combination of 5 µM rotenone (Sigma, 557368) and 5 µM antimycin A (Sigma, A8674).

Assay cycles consisted of 2 minutes of mixing followed by 3 minutes of measurement. The number of cycles performed was as follows: 6 for basal OCR, 4 for 20 mM glucose with 0.5mM glutamine and 3.5mM amino acid mix, 5 for oligomycin, 4 for FCCP, and 6 for rotenone + antimycin A.

#### Transplantation studies

Transplantation of SC-islets under the kidney capsule of 8-12 week-old immunodeficient SCID-beige mice was done as previously described.^9^ SC-islets were washed and resuspended in RPMI and submitted to University of Pennsylvania Stem Cell and Xenograft Core (SCXC – RRID: SCR_010035). The procedure was performed under anesthesia. Post-surgery, mice were monitored for pain and distress, with symptomatic mice isolated and observed for two days. If conditions did not improve, euthanasia was performed.

#### Insulinotropic agents screen

SC-islets were prepared and incubated in 2.8mM glucose KREBs buffer in typical static GSIS fashion. Insulinotropic agents were added to KREBs buffer during the high glucose (20 mM glucose) 1h incubation step at 37°C in a humidified 5% CO₂ incubator. The following compounds were used: dorzagliatin (10µM) (Selleckchem, S6921), forskolin (10 µM) (Stemolecule, 04-0025), PMA (10µM) (Santa Cruz Biotechnology, SC-3576B), ionomycin (10µM) (StemCell Technologies, 73722), and carbachol (1mM) (Santa Cruz Biotechnology, SC-202092). For select metabolic flux modulators, longer pre-incubation was required: cells were treated with either 1 mM dichloroacetate (DCA) (Selleckchem, S8615) or 1 mM oxamate (MedChemExpress, HY-W013032A) for 24 hours prior to GSIS. For oxamate-treated samples, 100µM methyl pyruvate (Sigma Aldrich, 371173) was also added during the 1-hour high-glucose stimulation step only. Following treatment, GSIS was performed as described above. Media samples were collected for insulin quantification, and data were normalized to cell counts.

#### Calcium influx

SC-islets were washed twice, followed by staining with 16.7μM Fura2-AM (Life Technologies; F-F1225) in Krebs buffer supplemented with 5mM glucose for 45 minutes at 37°C. Islets were then transferred to a perifusion chamber on a homeothermic platform, perifused with 37°C Krebs buffer at a flow rate of 1mL/min. and imaged with an inverted Zeiss microscope (Axio Observer.Z1). After imaging in the absence of substrate, a physiological amino acids mixture (0.5 mM glutamine and 3.5mM amino acid mix) alone and with increasing glucose levels (3mM, 16.7mM) were sequentially applied, followed by washing all substrates away and applying KCl (30mM). Intracellular Ca^2+^ was measured by dual-wavelength fluorescence microscopy using the Zeiss AxioVision system.

#### Key Resource Table

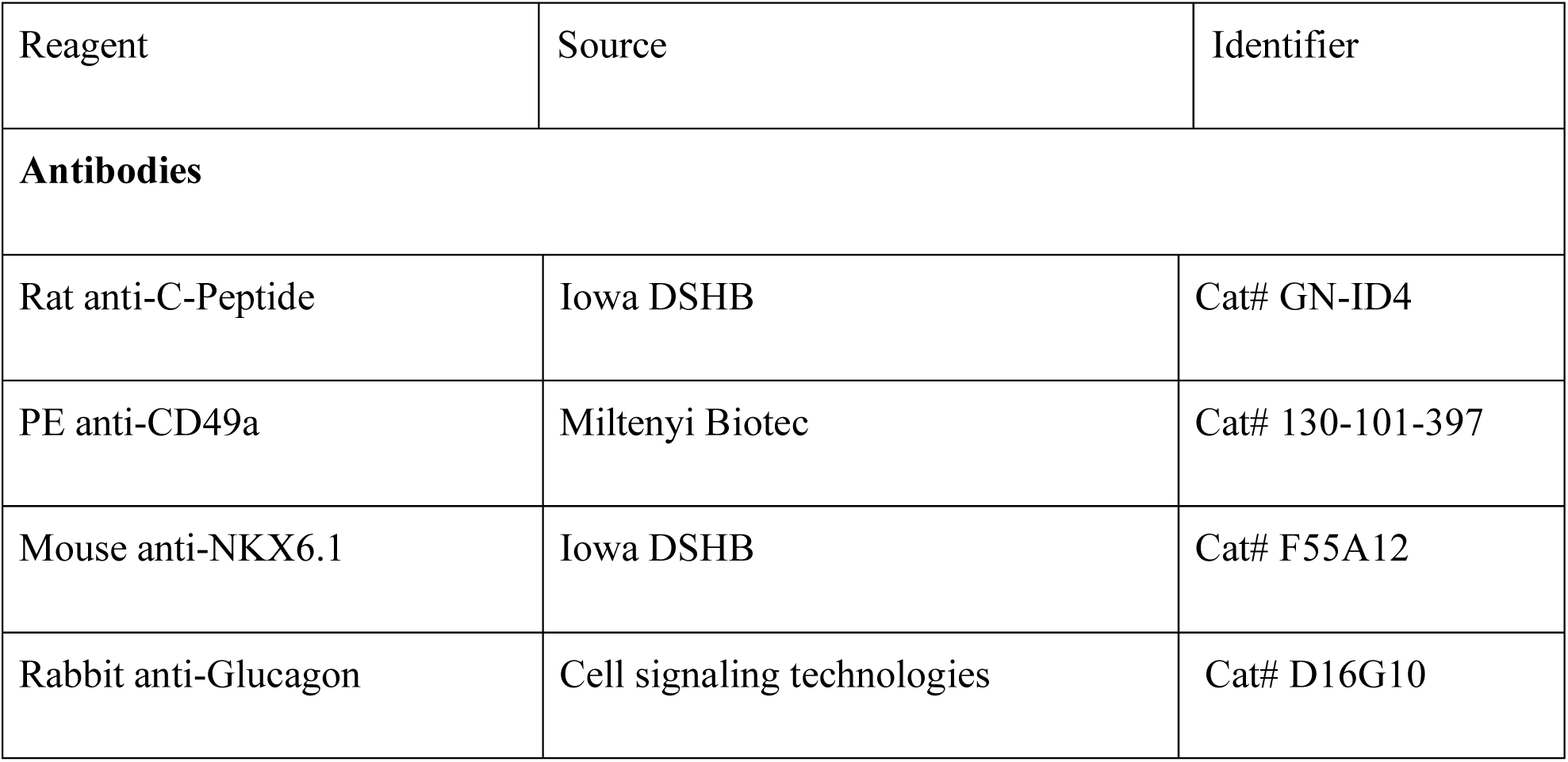
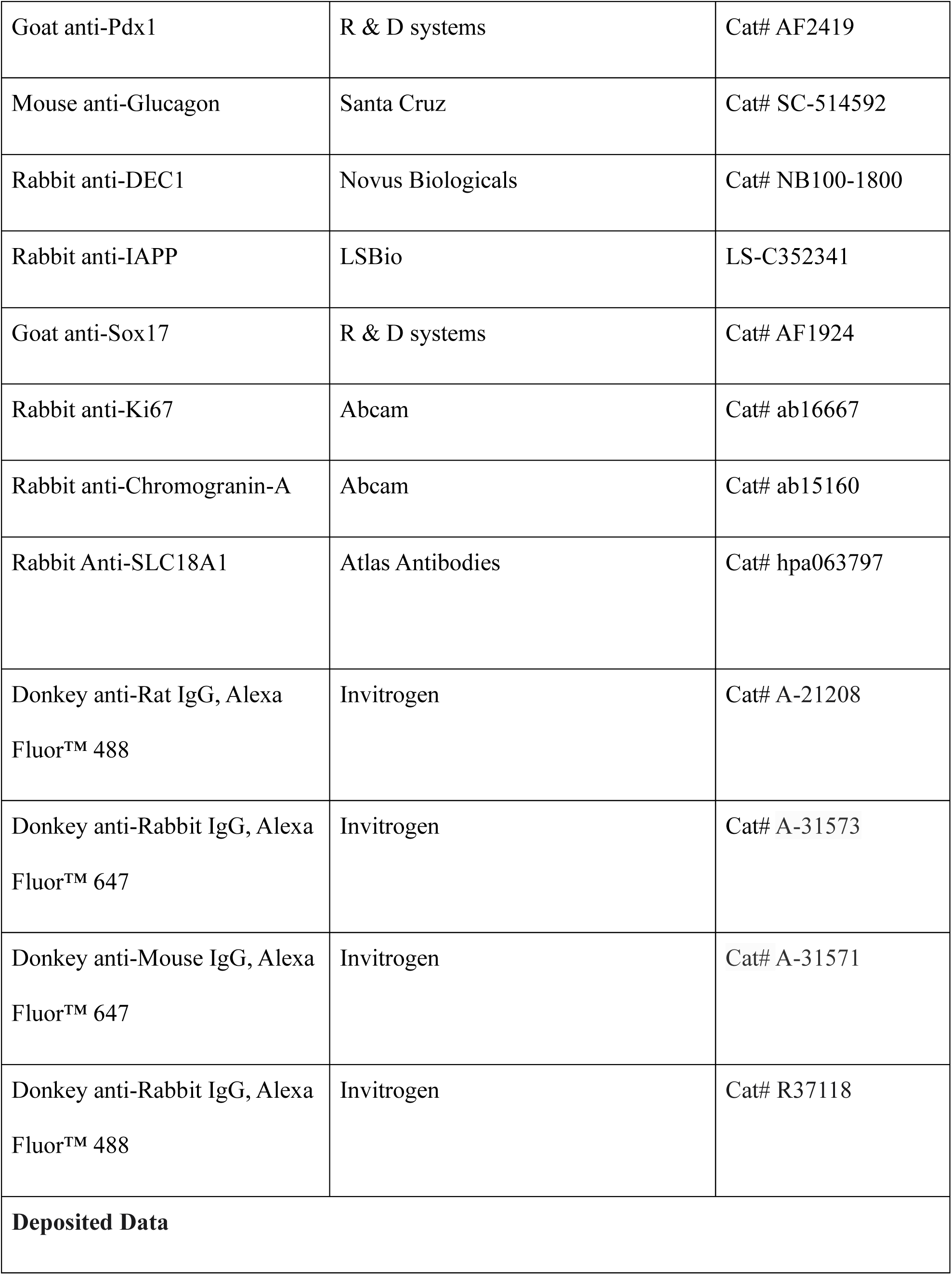
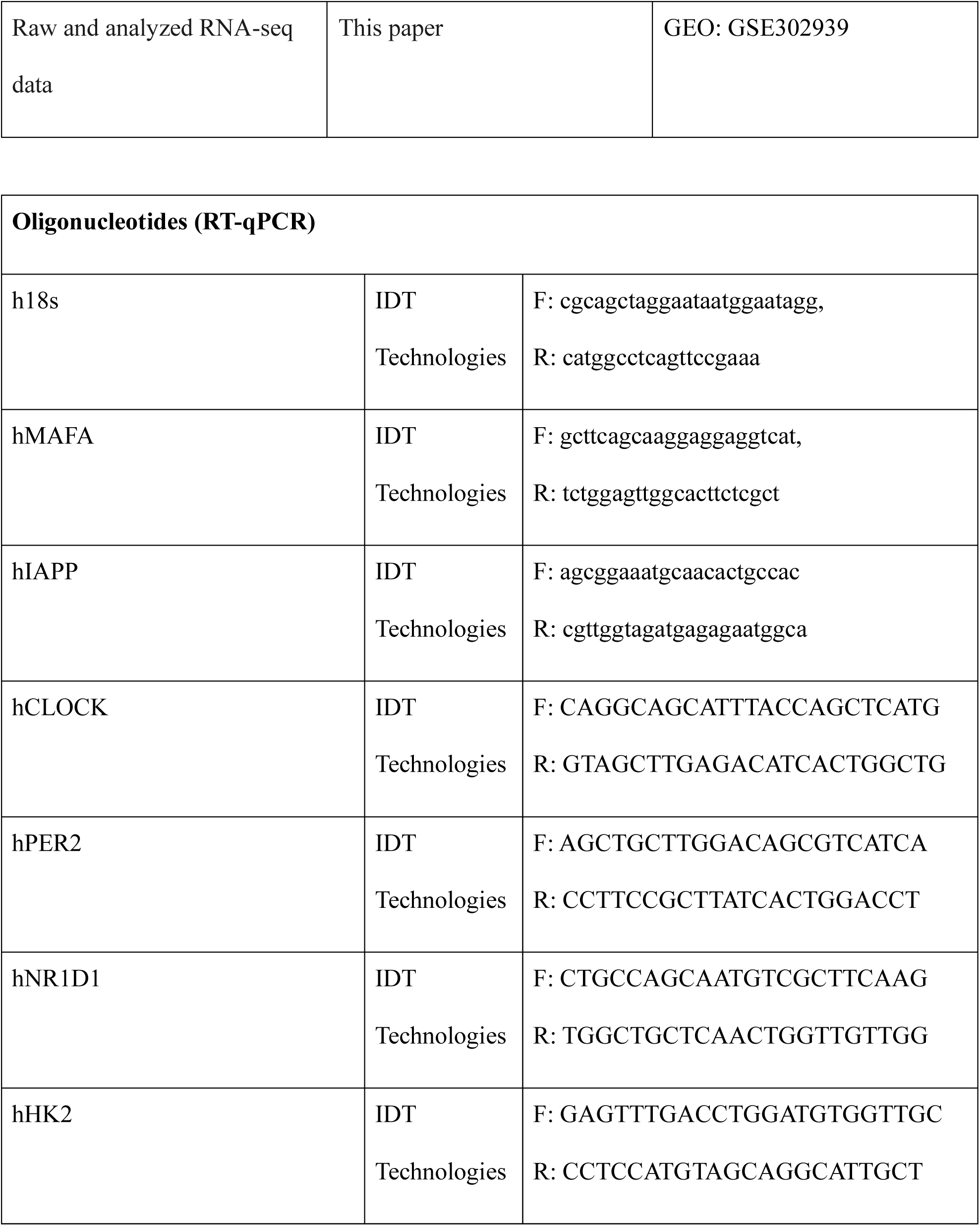
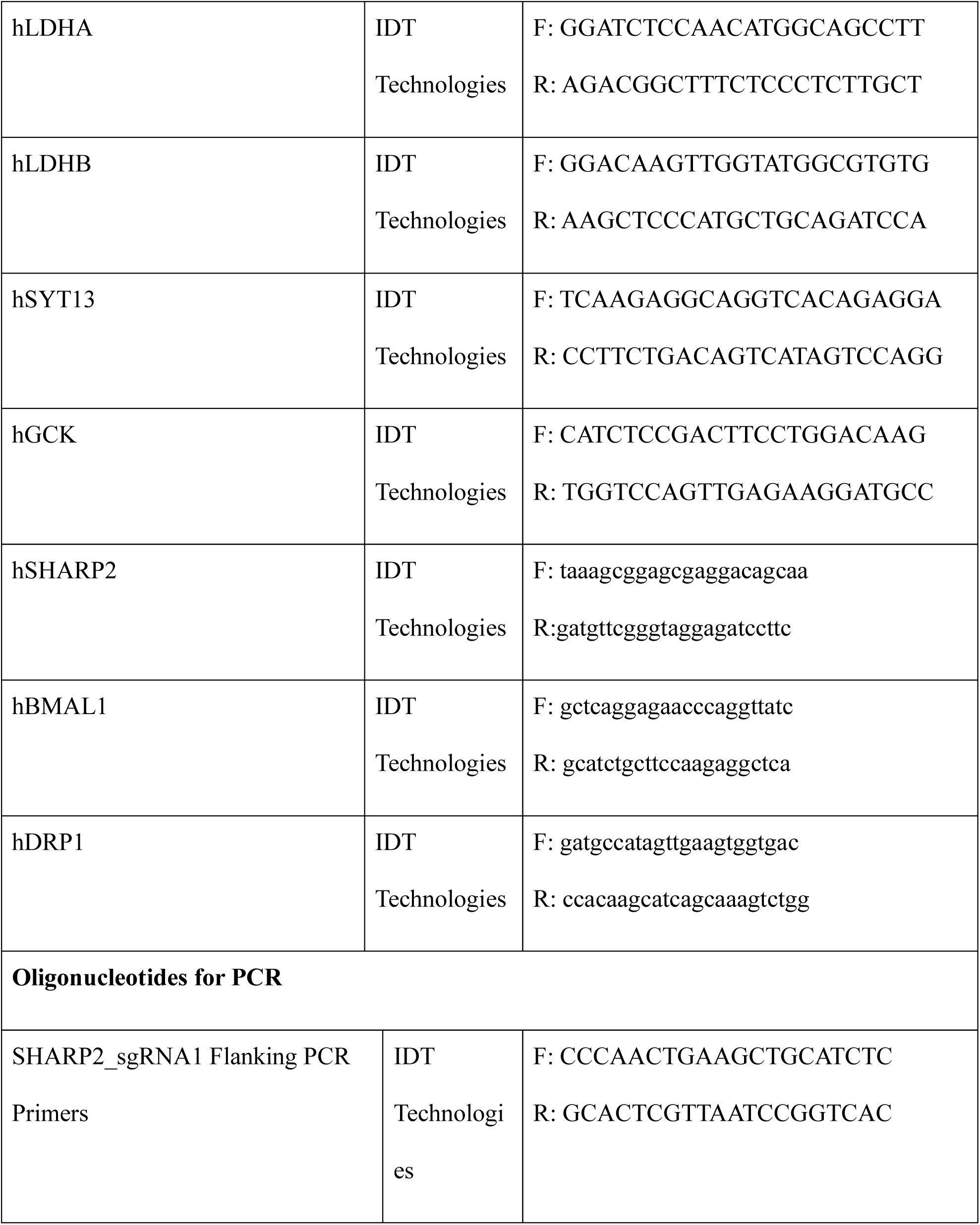
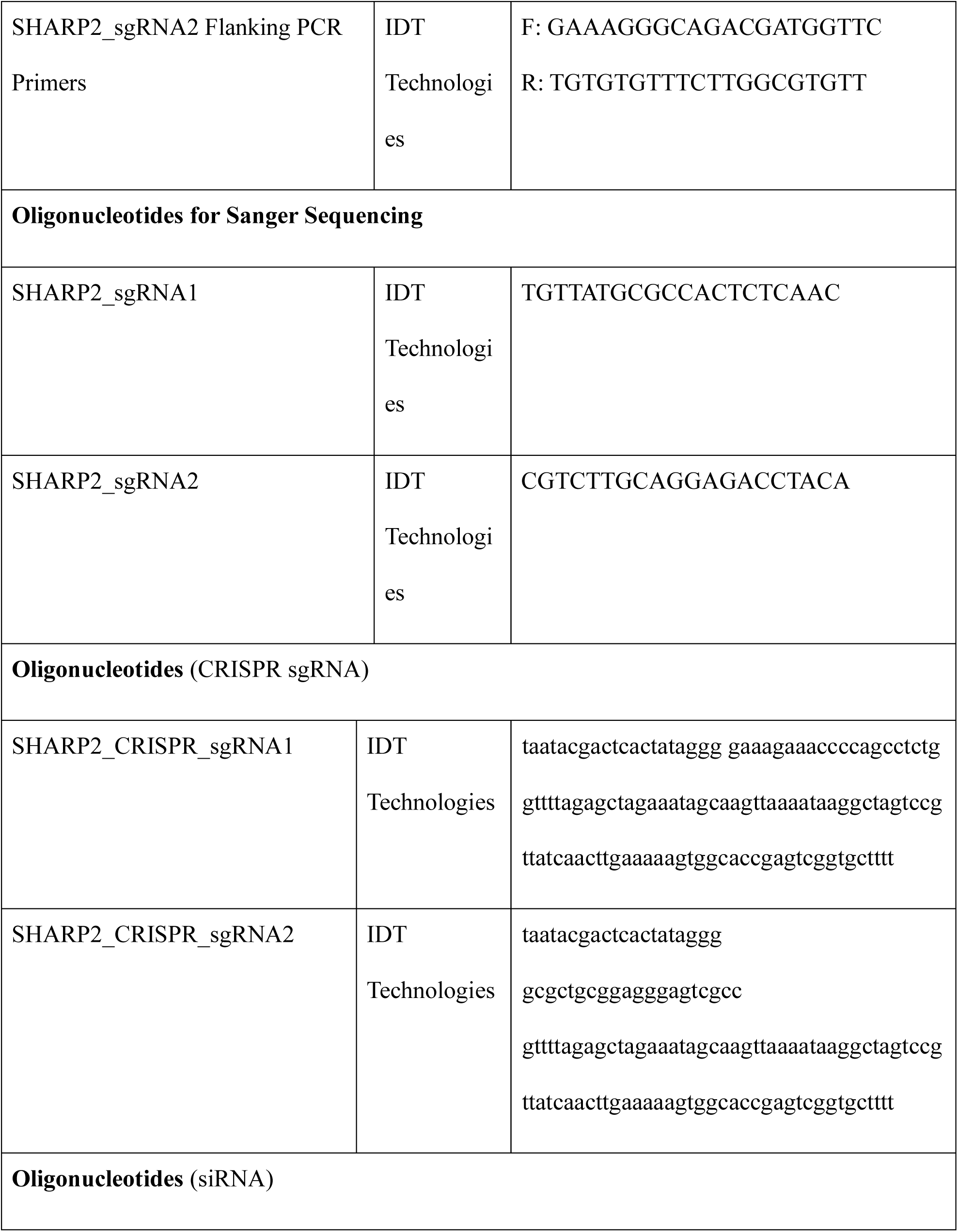
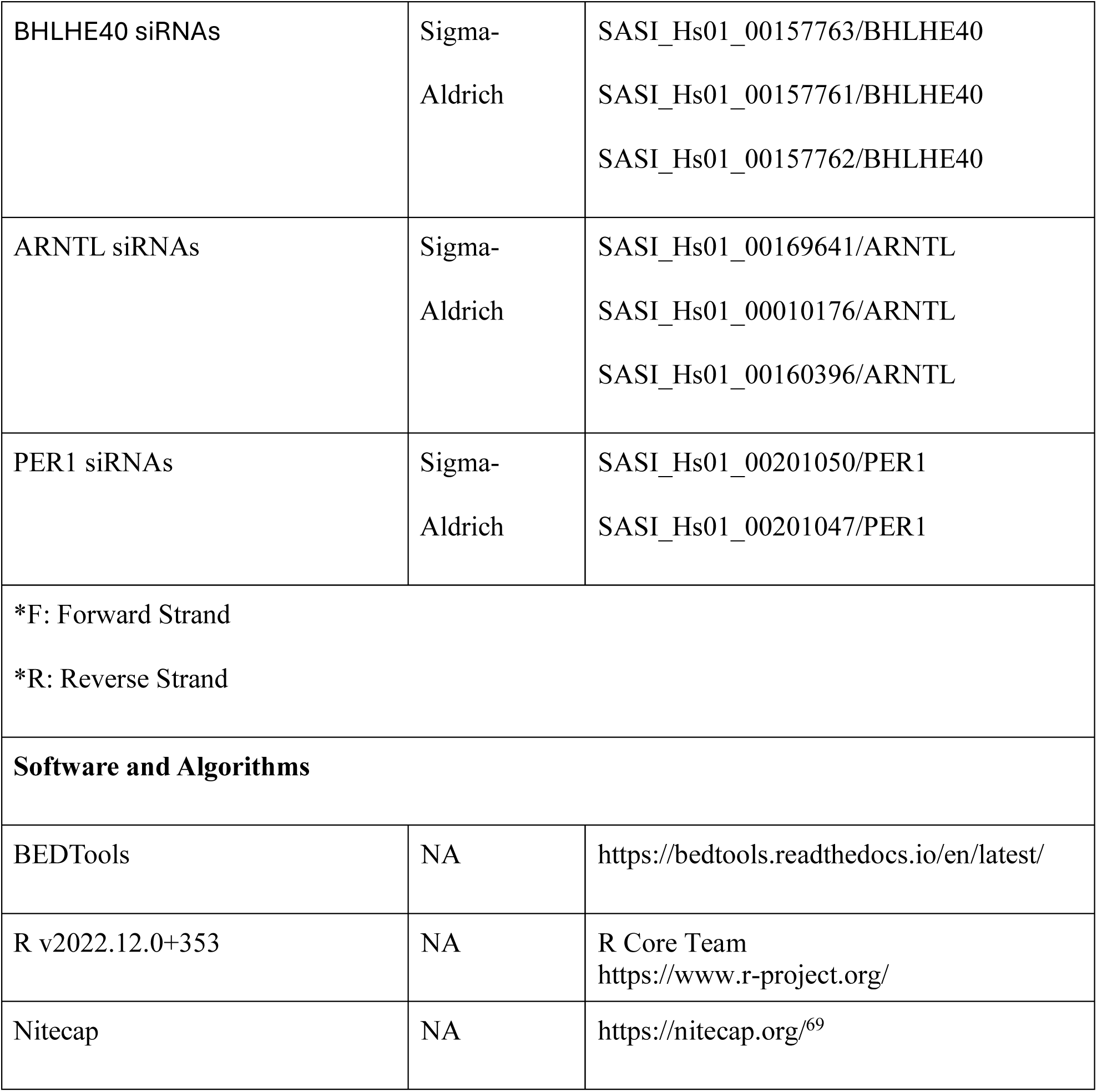

### QUANTIFICATION AND STATISTICAL ANALYSIS

No statistical methods were used to predetermine sample size or remove outliers. To assess the statistical difference between two sets of unpaired data, GraphPad Prism was used to calculate statistical significance for all non-RNA sequencing data. To assess the statistical difference between two sets of unpaired, the Shapiro-Wilk normality test was first performed as a column or grouped analysis. A two-sided unpaired t-test was then used to assess confidence on the measured difference of their mean values. For unpaired data that did not follow a normal distribution, we used a non-parametric Wilcoxon rank sum test to determine if they belong to the same parent distribution.

## EXCEL TABLE TITLES AND LEGENDS

**Table S1. SC-β cells differential gene expression output by DESeq. Related to Figure 2.**

**Table S2. Gene ontology from GSEA analysis. Related to Figure 2.**

**Table S3. SC-islets differential gene expression output by DESeq. Related to Figure S2.**

---

**Figure S1.**
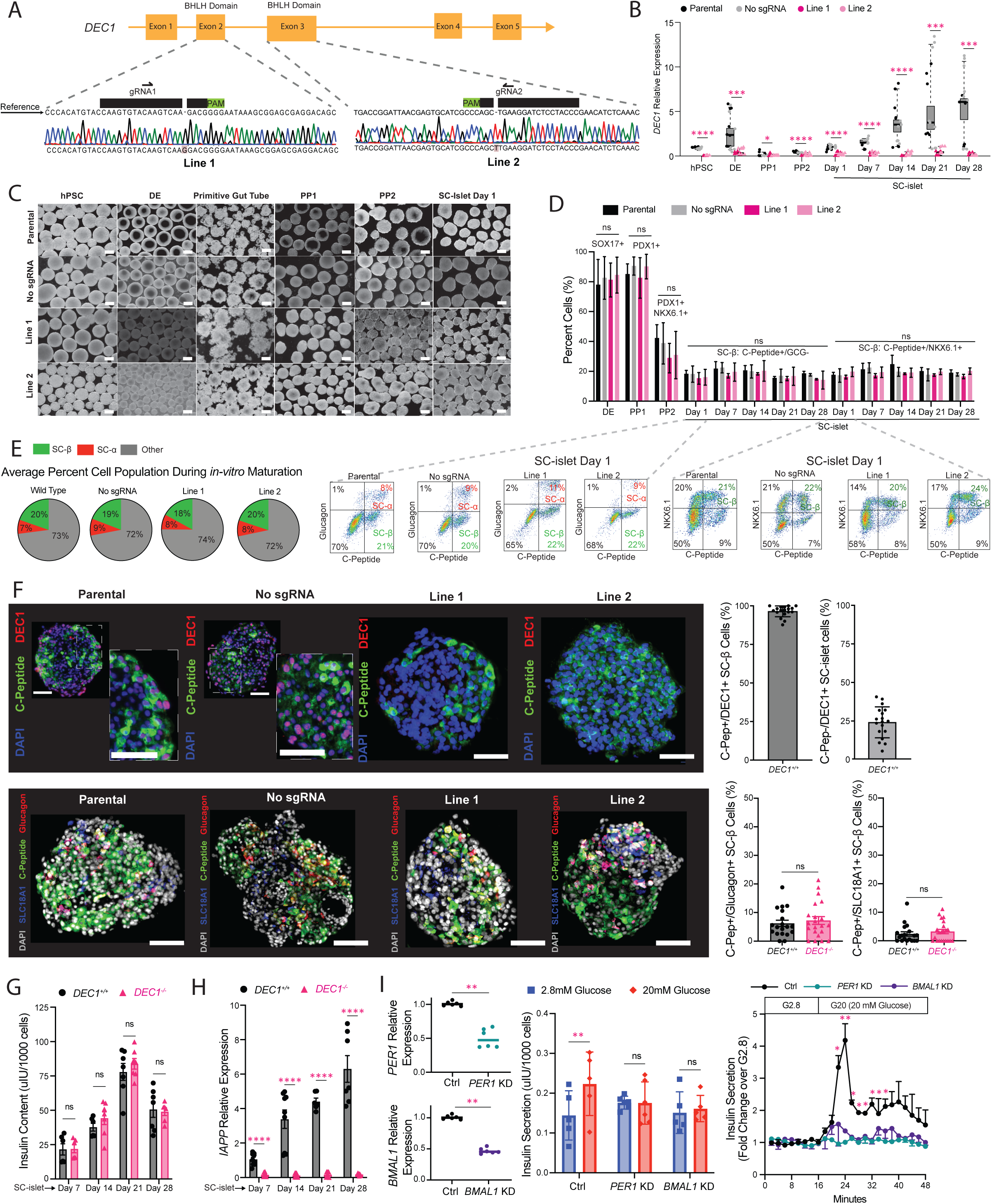
DEC1 has no effect on SC-islet differentiation, composition, or insulin production. Related to Figure 1. (A) Strategy to generate *DEC1*^-/-^ hPSC lines for subsequent differentiation. Shown are CRISPR/Cas9 sgRNAs used to create two *DEC1*^-/-^ hPSC lines, along with sequencing results indicating homozygous knockout due to frameshift-causing base-pair insertions. (B) *DEC1* expression is abrogated throughout *in vitro* differentiation and maturation of SC-islets generated from *DEC1*^-/-^ hPSC lines. Data from N = 4 differentiations with n = 2-3 technical replicate measurements. (C) 3D SC-islet organoids form normally from *DEC1*^-/-^ hPSC lines. Data are brightfield images of 3D organoids throughout differentiation. Scale bar is 100um. (D) SC-islets differentiate normally from *DEC1*-ablated hPSC lines. Data quantify the indicated cell populations by flow cytometry for the indicated differentiation markers (N = 5 differentiations, n ≥ 2000 cells each). Representative flow cytometry data are shown in S1E. (E) SC-islet composition is not affected by DEC1 loss. Data is average of SC-β (C-Peptide+/Glucagon-), and SC-α (Glucagon+) cells across day 7, 14, 21 and 28 for the indicated lines (N = 5 differentiations, n ≥ 2000 cells each). Representative flow cytometry data for differentiation marker quantification are shown to the right. (F) (Top) C-Peptide and DEC1 immunostaining of day 14 *DEC1^+/+^* and *DEC1^-/-^* SC-islets. Scale bar is 50um. Percent of C-peptide positive cells that are DEC positive, and percent of DEC1 positive cells that are C-peptide negative from N=5 *DEC1^+/+^* differentiations (n = 3-4 SC-islet organoids each) (right). (Bottom) Percent of polyhormonal cells is not affected by DEC1 loss. C-Peptide, Glucagon, and SLC18A1 immunostaining of day 14 *DEC1^+/+^* and *DEC1^-/-^* from N=5 *DEC1^+/+^* differentiations (n = 3-4 SC-islet organoids each). Percent of C-peptide positive cells that are glucagon positive, and percent of C-peptide positive cells that are SLC18A1 positive (right). (G) *DEC1*-ablated SC-islets produce insulin normally. Data is insulin content from day 7, 14, 21 and 28 *DEC1*^+/+^ and *DEC1*^-/-^ SC-islets (N=3 differentiation, n = 30-80 SC-islets each). (H) *IAPP* expression is significantly reduced in *DEC1^-/-^* SC-islets from week 1 through week 4 of extended *in vitro* culture. Data from N = 3 differentiation with n = 2-3 technical replicate measurements, normalized to day 7 SC-islets. (I) *BMAL1* and *PER1* siRNA knockdown (KD) in day 14 SC-islets show diminished glucose stimulated insulin secretion (GSIS) response. (Left) siRNAs successfully deplete *PER1* and *BMAL1* mRNA compared to siRNA control (Ctrl) conditions (N=3 differentiations, n=2 technical replicates each). (Middle) Diminished insulin response in static sequential incubations of 2.8mM glucose to 20mM glucose observed in day 14 *BMAL1* and *PER1* KD SC-islets (N=3 differentiations, n=50-100 islets each, assayed at least in duplicate). GSIS dynamics reveal day 14 *BMAL1* and *PER1* KD SC-islets sustain a low peak (first phase) insulin secretion under glucose stimulation (20mM) and dampened second phase secretion, consistent with the immature phenotype of *DEC1^-/-^*SC-islets. Data from day 14 *DEC1*^+/+^ (N=3 differentiation, n = 50-100 islets) perifused with the indicated substrates at least in duplicate, normalized to the mean of the first incubation. Data are mean ± SEM. *p <0.05, **p <1e-2, ***p <1e-3, ****p <1e-4, Wilcoxon test [(B), and (H)], and unpaired Welch’s t test [(D), (E) against Parental line; (F), (G), (H) and (I)].

**Figure S2.**
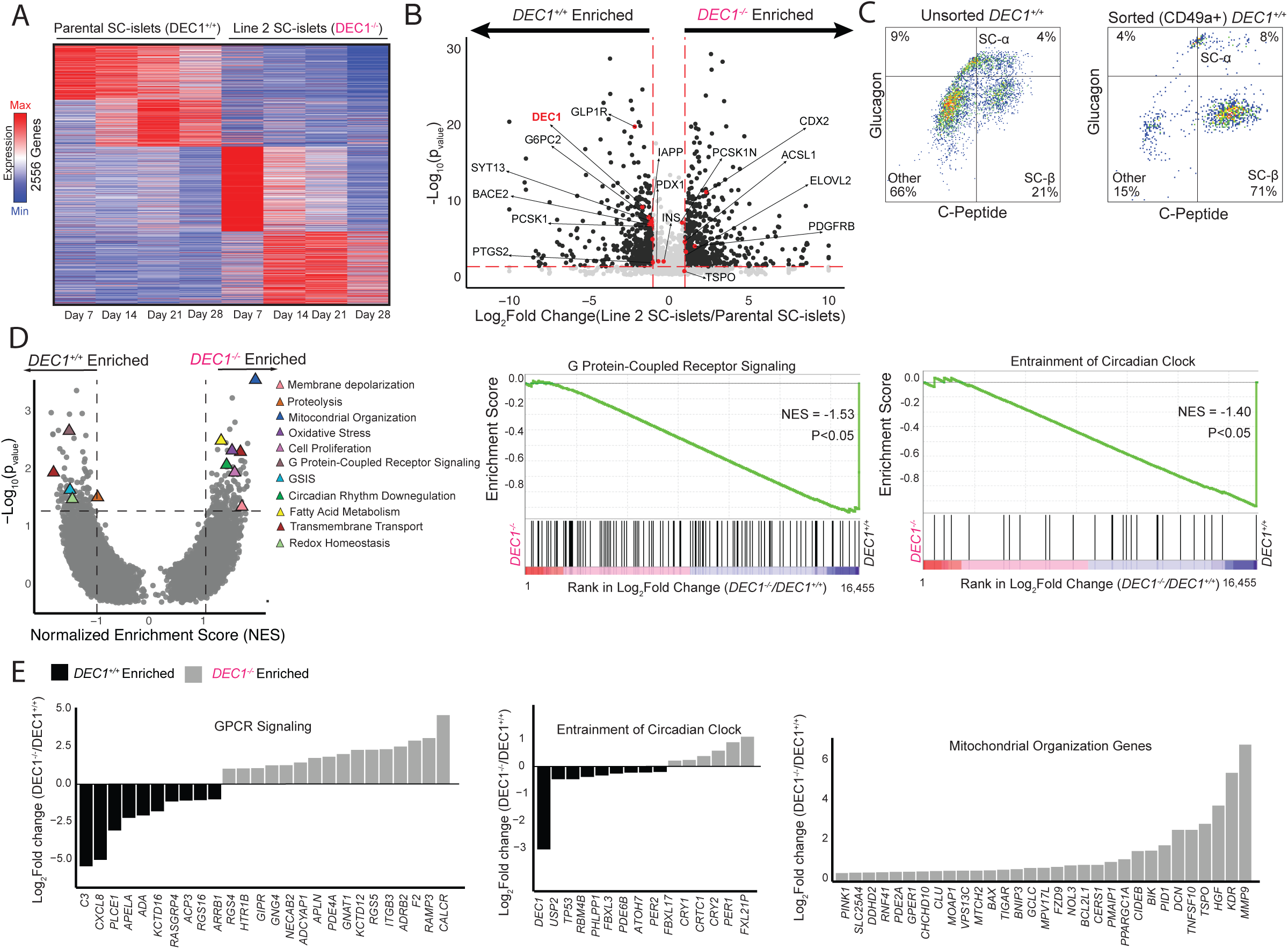
Additional gene set enrichment analysis of *DEC1^-/-^* SC-β cells. Related to Figure 2. (A) *DEC1* ablation in SC-islet organoids results in 2,556 differentially expressed genes. Heatmap shows differentially expressed genes (p < 0.05, DEseq test) in Parental and *DEC1^-/-^* Line 2 SC-islets at days 7, 14, 21, and 28 of *in vitro* maturation. (B) SC-β maturity-linked genes are depleted in *DEC1^-/-^* SC-islets. Volcano plot shows all differentially expressed genes at day 21 of SC-islet *in-vitro* maturation. Genes with at least a 2-fold change and p-value less than 0.05 are in black. Genes of interest are highlighted. (C) Flow cytometric sorting of CD49a-positive cells enables RNA-sequencing of highly enriched SC-β populations from day 14 SC-islets. n ≥ 2000 cells. (D) *DEC1* ablation results in dysregulation of important β-cell processes in SC-β cells (p-value < 0.05 and absolute NES value > 1) (left). Gene set enrichment analysis shows genes ranked by fold change in day 14 *DEC1^-/-^* vs. *DEC1^+/+^*SC-β cells (middle and right). NES, normalized enrichment score. (E) *DEC1* ablation upregulates genes modulating mitochondrial autophagy and mitochondrial membrane depolarization in SC-β cells. Shown are fold changes of genes in the indicated gene sets in day 14 SC-β cells.

**Figure S3.**
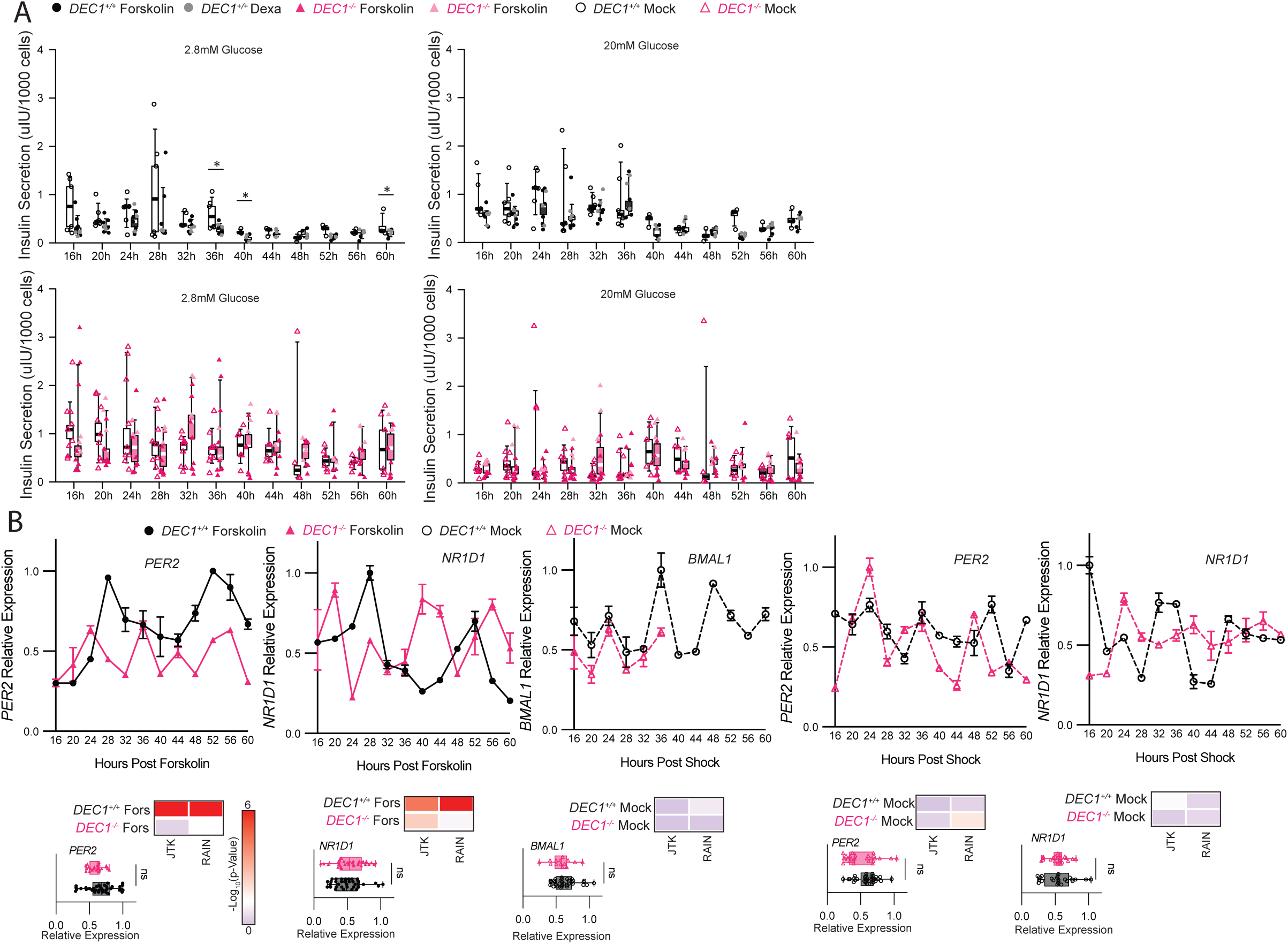
Circadian insulin secretion and clock gene expression in *DEC1^+/+^* and *DEC1^-/-^* SC-islets. Related to Figure 3. (A) GSIS responses show reduced insulin secretion in non-stimulatory (2.8mM) glucose at 36, 40, and 60 hours post forskolin or dexamethasone treatment in *DEC1^+/+^* SC-islets (top). Insulin secretion in 20mM glucose is shown to the right. Data are mean ± SEM. *p <0.05, unpaired Welch’s t-test. N=3 differentiations, with n=50-100 SC-islets assayed from each at least in duplicate. (B) *DEC1* loss disrupts circadian synchronization of the indicated genes. Data is expression relative to the maximum expression across the conditions shown in each panel (N = 3 differentiations with n =2 replicate measurements each). Rhythmicity p-values by JTK and RAIN analysis.

**Figure S4.**
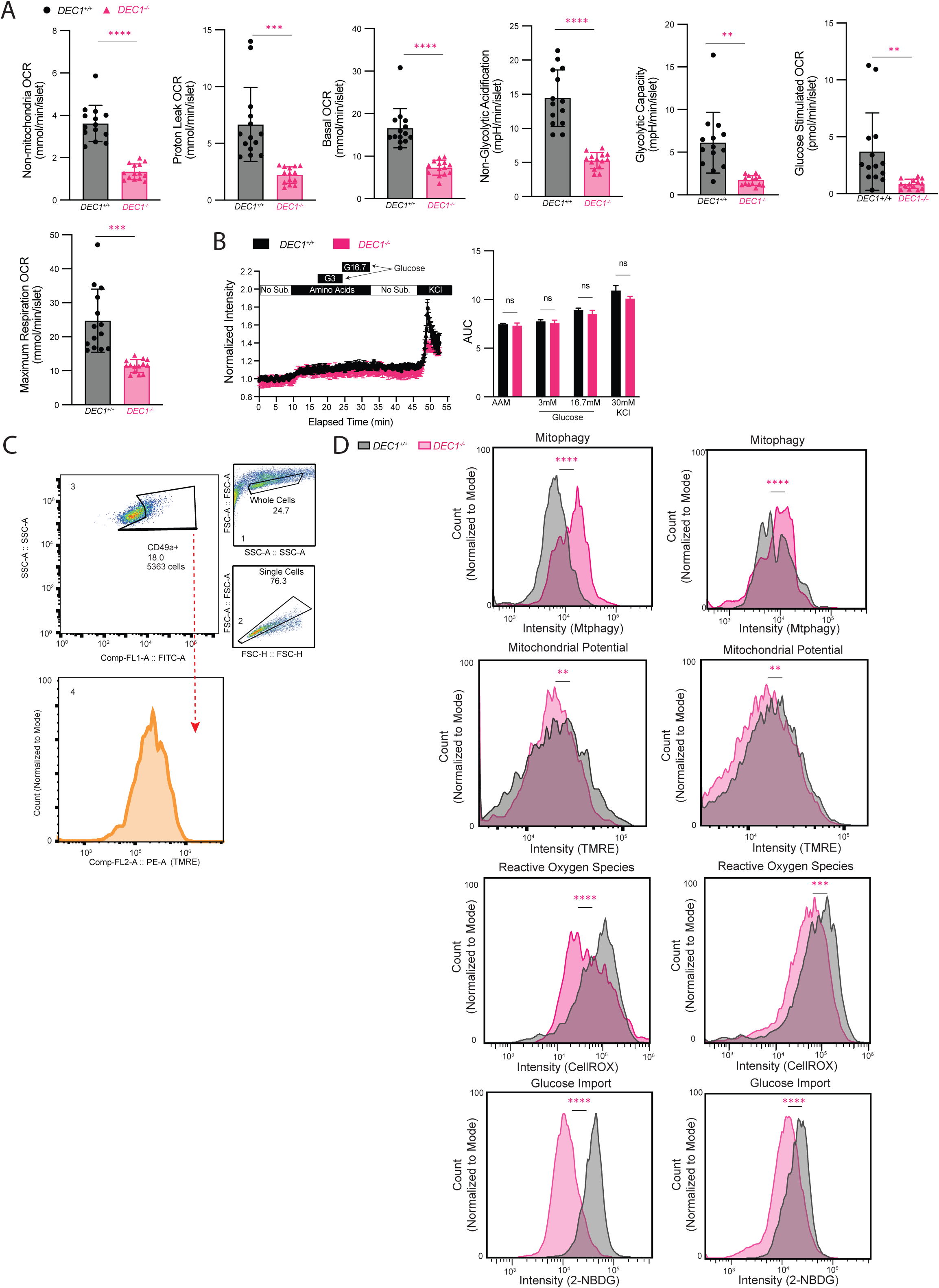
DEC1 regulates glycolytic and oxidative metabolism in maturing SC-β cells. Related to Figure 4. **(A)** Decreases in the indicated features determined by oxygen consumption rate (OCR) and extracellular acidification rate (ECAR) analysis of *DEC1^-/-^* SC-islets. Data from Figure 4B experiments (N = 4 differentiations, each with n = 3-4 replicate measurements of 8 similar-sized SC-islets). **(B)** Ca^2+^ influx measured by Fura-2 staining in response to no substrate (No Sub.); a physiological mixture of amino acids (AAM); AAM supplemented with 3mM glucose (G3); AAM supplemented with 16.7mM glucose (G16.7); no glucose; and 30 mM KCl, with respective AUCs quantified to the right. N=1 differentiation, n = 3 replicates with 8-12 similar-sized day 21 SC-islets. Data are mean ± SEM. *p <0.05, **p <1e-2, unpaired t-test. **(C)** CD49a staining allows for SC-β cell-specific staining of mitochondria integrity stains (TMRE, CellROX, and Mtphagy) and glucose analog reporter 2-NBDG. n = 5363 cells from day 14 SC-islets. **(D)** Related to Figure 4A. N=2 additional differentiations demonstrating *DEC1*-deficient day 14 SC-β cells exhibit mitochondria with increased autophagy and reduced membrane potential, along with lower reactive oxygen species generation and glucose import. n ≥ 2000 cells Data are mean ± SEM. *p <0.05, **p <1e-2, ***p <1e-3, ****p <1e-4, unpaired Welch’s t test [(A) and (B)] and Wilcoxon test (D).

